# A Polynomial-Time Algorithm for Minimizing the Deep Coalescence Cost for Level-1 Species Networks

**DOI:** 10.1101/2020.11.04.368845

**Authors:** Matthew LeMay, Ran Libeskind-Hadas, Yi-Chieh Wu

## Abstract

Phylogenetic analyses commonly assume that the species history can be represented as a tree. However, in the presence of hybridization, the species history is more accurately captured as a network. Despite several advances in modeling phylogenetic networks, there is no known polynomial-time algorithm for parsimoniously reconciling gene trees with species networks while accounting for incomplete lineage sorting. To address this issue, we present a polynomial-time algorithm for the case of level-1 networks, in which no hybrid species is the direct ancestor of another hybrid species. This work enables more efficient reconciliation of gene trees with species networks, which in turn, enables more efficient reconstruction of species networks.

## 1 Introduction

Reconstructing the evolutionary histories of a group of species is a fundamental step in phylogenetic analysis. While it is possible to infer trees from whole-genome alignments or from concatenated alignments, a common approach relies on first reconstructing individual *gene trees*, then reconstructing a *species tree* from the gene trees. However, gene trees and species trees may be incongruent due to various evolutionary processes, thus requiring *reconciliation* methods that map a gene tree “within” a species tree and explain topological differences by postulating a sequence of evolutionary events, with different models allowing for different types of events.

In the popular *multispecies coalescent (MSC) model* (Maddison 1997), species are treated as populations of individuals, and incongruence is assumed to be caused by *incomplete lineage sorting (ILS)* (Figure 1a,b). Formally, two lineages may fail to coalescence at their most recent opportunity, a phenomenon known as *deep coalescence*. ILS occurs when one lineage then coalesces with a lineage from a less closely-related population (Degnan and Rosenberg 2009).

**Figure 1.**
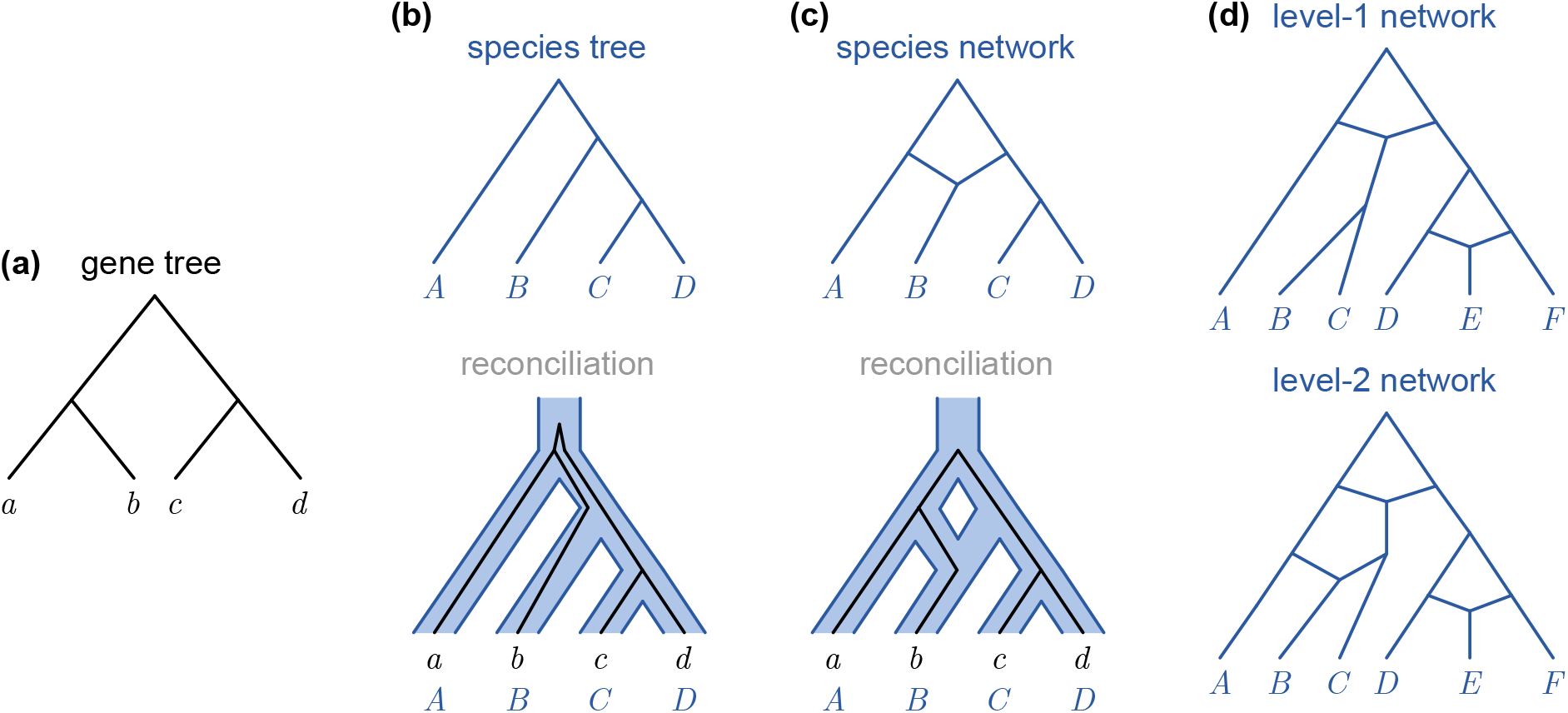
Gene trees, species trees, and species networks. (a) A gene tree. (b) A species tree and reconciliation. Under the multispecies coalescent model, the gene tree evolves within the species tree, and incongruence between the trees is due to ILS. (c) A species network and reconciliation. The same gene tree evolves within the species network, and no ILS is necessary. (d) A level-1 species network and a level-2 species network.

Coalescent theory allows for computing the probability of a gene tree topology given a species tree topology and parameters such as population size and divergence time (Kingman 1982, Wakeley 2008). Thus, given multiple gene trees, it is possible to infer a species tree using either probabilistic or parsimony approaches (see Degnan and Rosenberg (2009) for a review of such methods). Probabilistic approaches rely on maximum likelihood or Bayesian estimation, whereas a parsimony approach chooses a species tree by minimizing deep coalescences (MDC), which “minimizes the number of extra lineages that had to coexist along species lineages” (Maddison 1997). In general, parsimony approaches require only topologies and are more efficient than probabilistic approaches, and thus are more broadly applicable.

However, the MSC model commonly assumes that species histories can be represented as a tree and therefore cannot account for hybridization (Figure 1c), in which separate species exchange genetic information, either through introgression, in-breeding between separate populations, or hybrid speciation (Elworth et al. 2019, Runemark et al. 2019). Studies have shown that hybridization can play a role in the evolution of eukaryotic species (Mavárez et al. 2006, Fontaine et al. 2015, Racimo et al. 2015, Lamichhaney et al. 2018).

In the last decade, several algorithms have been developed to infer species networks by simultaneously modelling ILS and hybridization. In a species network, species branches can join together at *hybridization nodes* (also known as *reticulation nodes*). As with the simpler MSC model, there exist both probabilistic (Meng and Kubatko 2009, Kubatko 2009, Yu et al. 2011a; 2012; 2013b; 2014, Wen and Nakhleh 2017, Zhang et al. 2017) and parsimony approaches (Yu et al. 2011a; 2013a;b) for inferring species networks under these models. Many of the parsimony approaches rely on converting a species network to a multi-labeled tree (MUL-tree), then considering all mappings of alleles to their sampled species. Because there can exist an exponential number of allele mappings, such approaches may not scale to large numbers of species or hybridizations.

Rather than model ILS and hybridization, some models instead allow for ILS and horizontal gene transfer, often with gene duplication and loss (Stolzer et al. 2012, Chan et al. 2017). However, such models also assume the species history can be represented as a tree and that gene transfers result in gene trees that are incongruent with the species tree. In contrast, by relying on a species network rather than a tree, hybridization allows different segments of the gene tree to have different histories naturally by using different edges leading to a hybridization node.

In parallel with these advances in ILS and hybridization, To and Scornavacca (2015) developed two algorithms for reconciling gene trees and species networks that take into account duplication and loss events. They studied two variations: first, finding an optimal tree in a network such that the reconciliation of the gene tree and the “displayed” species tree has minimum cost, and second, finding a minimum cost reconciliation between the gene tree and the full species network. Interestingly, the time complexity of their first algorithm depends not on the number of hybridization events but on a parameter of the network called its *level* (Choy et al. 2005a), intuitively a measure of “how much the network is ‘tangled”‘ (To and Scornavacca 2015) or how densely its hybridization nodes are distributed (Figure 1d). This algorithm is fixed-parameter tractable when parameterized by the level of the network and the number of biconnected components in the network. Their second algorithm is polynomial in the number of hybridization nodes, the size of the gene tree, and the size of the species network.

Despite these advances, there is currently no known polynomial-time algorithm for inferring an MDC reconciliation between a gene tree *G* and a species network *S*. We address this challenge with the following contributions:

1. We present a O(|*G*|·|*S*|^5^) algorithm for reconciling a gene tree *G* and species network *S* when *S* has one hybridization node. Like many parsimony approaches, our algorithm relies on dynamic programming. Our key insight is to introduce a new parameter of the reconciliation, the *signature*, which specifies which hybridization edges are used by different parts of the reconciliation.
2. We reduce the time complexity of the previous algorithm to O(|*G*| · |*S*|^2^) by generalizing the concept of a single lowest common ancestor (LCA) in trees to multiple LCAs in networks.
3. We present a O(|*G*| · |*S*|^2^) algorithm for reconciling *G* and *S* when *S* is a level-1 network. Intuitively, in a level-1 network, no hybrid species is the direct ancestor of another hybrid species. For a general level-*k* network, the time complexity increases to O(4^*k*^ · |*G*| · |*S*|^2^), which, while exponential, is still smaller than existing algorithms that are exponential in the number of species and hybridization nodes.

## 2 Background

### 2.1 Preliminaries

We start by giving some basic definitions using notation largely verbatim from To and Scornavacca (2015). A summary of notation can be found in Supplemental Table S1.1.

A *rooted phylogenetic network* refers to a rooted directed acyclic graph with a single root with in-degree 0 and out-degree 2, multiple leaves with in-degree 1 out-degree 0, and internal nodes with either in-degree 1 and out-degree 2, or in-degree 1 and out-degree 2. Nodes in this last set are *hybridization nodes*, and edges leading to hybridization nodes are *hybridization edges*. Given a network *N*, let *V*(*N*) denote its node set and *E*(*N*) denote its edge set. Let *L*(*N*) ⊂ *V*(*N*) denote its leaf set, *I*(*N*) = *V*(*N*) \ *L*(*N*) denote its set of internal nodes, and *r*(*N*) ∈ *I*(*N*) denote its root node. For node *v* ∈ *V*(*N*), let *c*(*v*) denote its set of children, *p*(*v*) denote its parent (either a single node or a set of two nodes), and, if *v* has a single parent, *e*(*v*) denote the edge (*p*(*v*),*v*). The size of *N*, denoted by |*N*|, is equal to |*V*(*N*)| + |*E*(*N*)|. Given *v* ∈ *V*(*N*), let *N_v_* denote the subnetwork of *N* rooted at *v*, i.e. the subgraph of *N* consisting of all nodes and edges reachable from *v*.

Define ≤_*N*_ (<_*N*_) to be the partial order on *V*(*N*), where given two nodes *u* and *v* of *N*, *u* ≤_*N*_ *v* (*u* <_*N*_ *v*) if and only if there exists a path in *N* from *v* to *u* (and *u* ≠ *v*). The partial order ≥_*N*_ (>_*N*_) is defined analogously. In such a case, *u* is said to be *lower or equal to* (lower than) *v*, and *u* a (strict) *descendant* of *v*, and *v* a (strict) *ancestor* of *u*.

Given two nodes *u* and *v* of *N* such that *u* ≤_*N*_ *v*, a path from *v* to *u* in *N* is a sequence of contiguous edges from *v* to *u* in *N*. Note that if *v* = *u*, the path from *v* to *u* is empty. As there can be multiple paths between pairs of vertices in a network, let *paths_N_*(*v, u*) denote the set of all paths from *v* to *u*. Let *paths*(*N*) denote the set of all paths in network *N*.

Let 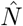 be the underlying undirected graph corresponding to *N*. An undirected graph is said to be biconnected if it remains connected when any single node is removed. A subgraph of a graph 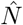 is said to be a *biconnected component* if it is a maximal biconnected subgraph of 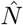. If every biconnected component of 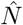 has at most *k* hybridization nodes, we say that *N* is of level-*k* (Choy et al. 2005b). A *rooted phylogenetic tree* is a rooted phylogenetic network with no hybridization nodes, i.e. a level-0 network. In the remainder of this paper, we refer to rooted phylogenetic networks and rooted phylogenetic trees simply as *networks* and *trees*, respectively. We allow trees to contain *artificial nodes*, i.e. nodes with in-degree and out-degree 1, and *handle nodes*, i.e. nodes with in-degree 0 and out-degree 1.

Let a *species network S* depict the evolutionary history of a set of species, and let a *gene tree G* depict the evolutionary history of a set of genes sampled from these species. To compare a gene tree with a species network, let a *leaf map Le*: *L*(*G*) → *L*(*S*) label each leaf of the gene tree with the leaf of the species network from which the gene was sampled. The mapping need not be one-to-one nor onto.

### 2.2 Reconciliations

#### Definition 2.1

(Reconciliation). Given a gene tree *G*, a species network *S*, and a leaf map *Le*, a *reconciliation*^1^ *R* for (*G, S, Le*) is a pair of mappings (*R_v_, R_p_*) where *R_v_*: *V*(*G*) → *V*(*S*) is a *vertex mapping* and *R_p_*: *V*(*G*) → *paths*(*S*) is a *path mapping* subject to the following constraints:

1. If *g* ∈ *L*(*G*), then *R_v_*(*g*) = *Le*(*g*).
2. If *g* ∈ *I*(*G*), then for each *g*′ ∈ *c*(*g*), *R_v_*(*g*′) ≤_*S*_ *R_v_*(*g*).
3. If *g* ≠ *r*(*G*), then *R_p_*(*g*) ∈ *paths_S_*(*R_v_*(*p*(*g*)), *R_v_*(*g*)). Otherwise, *R_p_*(*g*) = 0.

Constraint 1 asserts that *R_v_* extends the leaf map *Le*. Constraint 2 asserts that *R_v_* satisfies the temporal constraints implied by *S*. Constraint 3 asserts that the vertex mapping and path mapping are consistent.

The vertex mapping specifies which node of *S* a node of *G* is mapped to, and the path mapping specifies a path in *S* between a node of *G* and its parent. Note that if *S* is a tree, a reconciliation can be specified by the vertex mapping alone, and the paths between nodes in the species tree would be implied. However, when hybridization is allowed, there can exist multiple paths between nodes in the species network, thus requiring the path mapping.

It will be convenient to consider several variants of a reconciliation. In the first, given *g* ∈ *V*(*G*), a reconciliation *R^g^* denotes the reconciliation *R* restricted to subtree *G_g_*. In the second, a reconciliation is restricted to a subnetwork of the species network (that consists of a subset of nodes and all edges between those nodes), and only the parts of the gene tree that evolve within the sub-network are considered. In the third, a reconciliation is extended to a forest of multiple gene trees, all of which evolve within the same species network. Henceforth, the term reconciliation encompasses these variants.

As typical in a multispecies coalescent process, evolution in the species network is viewed backward in time, from the leaves toward the root. Then, given a reconciliation *R*, one can directly count the number of gene lineages “exiting” each edge *e* of the species network. Specifically, given edge *e* ∈ *E*(*S*),

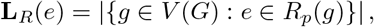

and the number of extra lineages is

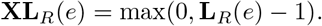

Finally, the *deep coalescence cost* of a reconciliation is the sum of extra lineages across all edges of the species network:

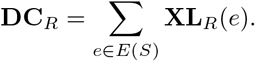

This value is also known as the *reconciliation cost*. Given a reconciliation *R*, the *edgeset* of *R* is the set of species edges used in the path mapping:

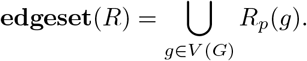

Clearly, for *e* ∈ **edgeset**(*R*), **XL**_*R*_(*e*) = **L**_*R*_(*e*) − 1, and thus, the following is an equivalent definition for the deep coalescence cost:

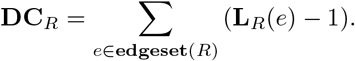

Finally, we define the Most Parsimonious Reconciliation Problem:

#### Problem 2.1

(Most Parsimonious Reconciliation (MPR)). Given *G, S*, and *Le*, the *MPR problem* is to find a reconciliation with minimum cost.

When *S* is a tree, the MPR is unique (the LCA reconciliation^2^) (Wu and Zhang 2011) and can be found in *O*(|*G*| · |*S*|) time (Zmasek and Eddy 2001). However, when *S* is a network, the MPR is not necessarily unique.

In this work, we consider the MPR Problem for special case of a binary gene tree and a level-1 species network.

## 3 Methods

In this section, we provide a polynomial-time algorithm for inferring an MPR between a binary gene tree *G* and a level-1 species network *S* with leaf map *Le*. Like many parsimony approaches, our algorithm relies on dynamic programming. For the sake of simplicity, we present only the algorithm for minimizing the cost of a reconciliation. By using standard annotation of the dynamic programming table, we can subsequently perform a traceback and reconstruct an actual reconciliation. A summary of notation can be found in Supplemental Table S1.2. For brevity, proofs appear in Supplemental Section S2.

In the remainder of this section, also for the sake of simplicity, we often omit *Le* in our exposition and theorems, with the understanding that given a gene tree *G* and species network *S*, we are given a leaf map *Le* as well. We do make the dependence on *Le* explicit in our algorithms.

### 3.1 Starting with a Simpler Problem

We start with the simpler problem of reconciling a gene tree *G* to a species network *S* with a single hybridization node *v^H^*. For two nodes *u* and *v* of *S*, it can be easily shown that there exists a single node, called the *lowest common ancestor (LCA)* and denoted *lca_S_*(*u, v*), that is the lowest element of *S* that is an ancestor of both *u* and *v*. Let *v^L^* and *v^R^* denote the left and right parents of *v^H^*, and let *e^L^* = (*v^L^, v^H^*) and *e^R^* = (*v^R^, v^H^*) denote the left and right edges to *v^H^*. Let *v^A^* = *lca_S_*(*v^L^, v^R^*), called the *split node*, denote the lowest common ancestor of *v^L^* and *v^R^* (Figure 2a).

**Figure 2.**
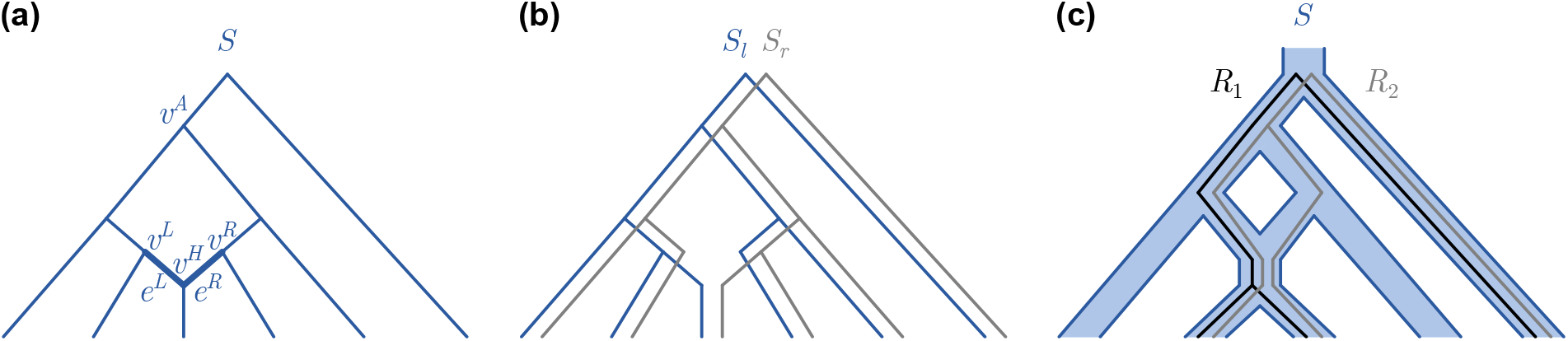
Species networks with one hybridization node. (a) Important nodes are labeled, including the hybridization node *v^H^*, the left and right parents *v^L^* and *v^R^* of *v^H^*, the split node *v^A^*, and the left and right edges *e^L^* and *e^R^* to *v^H^*. (b) Tree *S_l_* constructed from *S* with *e^L^* removed, and tree *S_r_* constructed from *S* with *e^R^* removed. (c) Two reconciliations *R*_1_ and *R*_2_ between the same gene tree and species tree. *R*_1_ subsumes *R*_2_.

Let 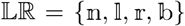 be a set of four symbols which will be used to denote the hybridization edges in a set. Because the species network has a single hybridization node, there are four options: none, left edge, right edge, both edges. For a set of elements *X*, let 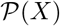 denote the power set over the elements. Then, for a set of edges *E* ∈ *E*(*S*), define a function **signature**(*E*): 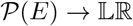 that denotes the hybridization edges in the set:

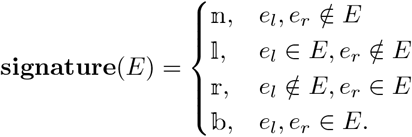

Define the binary operator + over 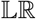 as follows:

- For each 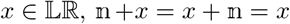.
- For each 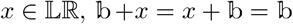.
- 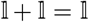 and 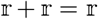.
- 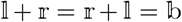.

It is easily verified that given two subsets *E*_1_ and *E*_2_ of *E*(*S*),

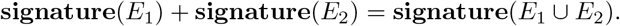

Given a reconciliation *R*, the signature of *R*, denoted **signature**(*R*), is defined to be the signature of **edgeset** (*R*). Conceptually, the signature of a reconciliation *R* denotes whether *R* uses neither edge, only the left edge, only the right edge, or both edges leading to the hybridization node.

#### Lemma 3.1

(Equivalent Edgesets). *Given a gene tree G and a species network S with one hybridization node, let* 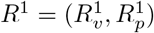 *and* 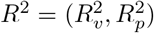 *denote two reconciliations between G and S. If* 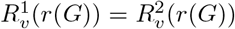 *and* **signature**(*R*^1^) = **signature**(*R*^2^), *then* **edgeset**(*R*^1^) = **edgeset**(*R*^2^).

Given a gene tree *G* and species network *S*, recall that a reconciliation is said to be *optimal* if it has the minimum cost among all reconciliations between *G* and *S*. A reconciliation *R* between *G* and *S* is said to be *root-signature-optimal (rs-optimal)* if it has the minimum cost among all reconciliations *R*′ between *G* and *S* such that 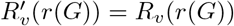 and **signature**(*R*′) = **signature**(*R*).

**Algorithm 1.**
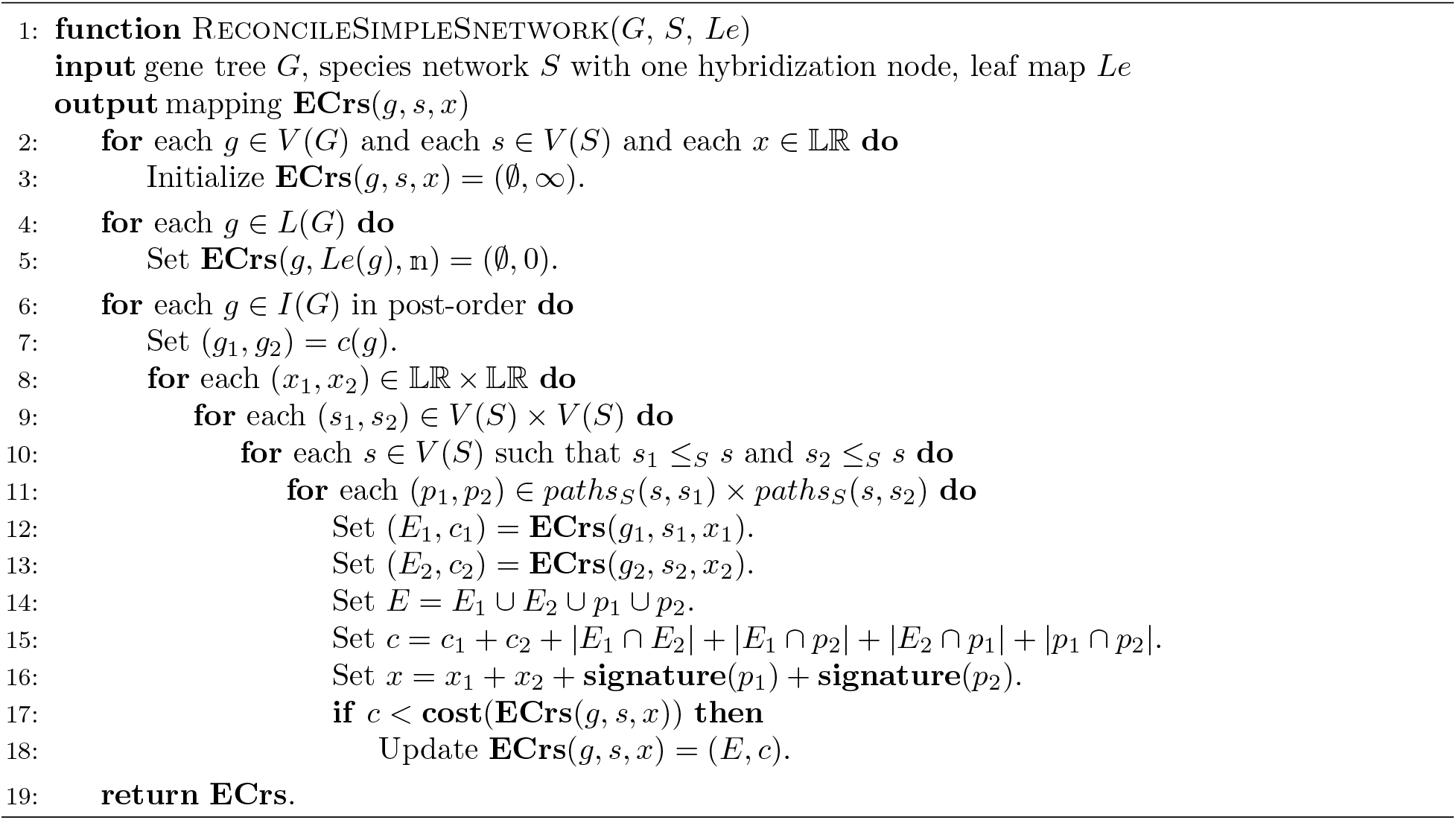

#### Lemma 3.2

(Optimal Substructure Property). *Given a gene tree G and a species network S with one hybridization node, let* 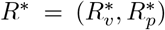 *be an rs-optimal reconciliation between G and S. Then for each g* ∈ *V*(*G*), *R*^*,*g*^, *the reconciliation R*\* *restricted to G_g_, is rs-optimal*.

We are now ready to describe our dynamic programming algorithm for reconciling *G* and *S*. Our algorithm constructs a dynamic programming table **ECrs**, where given any *g* ∈ *V*(*G*), *s* ∈ *V*(*S*), and 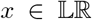, entry **ECrs**(*g, s, x*) is an ordered pair (*E, c*) for an rs-optimal reconciliation *R* = (*R_v_, R_p_*) between *G_g_* and *S* such that *R_v_*(*g*) = *s* and **signature**(*R*) = *x*. 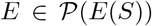 denotes the edgeset of *R*, and non-negative integer *c* denotes the cost of *R*. Note that by Lemma 3.1 and the definition of rs-optimality, the specific vertex mapping *R_v_* and path mapping *R_p_* do not matter: all rs-optimal reconciliations such that *R_v_*(*g*) = *s* and **signature**(*R*) = *x* share the same edgeset and cost. By Definition 2.1, if *g* ∈ *L*(*G*), then *R_v_*(*g*) = *Le*(*g*) and *R_p_*(*g*) = 0. In this base case, the reconciliation uses neither of the two hybridization edges and has cost 0. Otherwise, the algorithm considers *g* ∈ *I*(*G*) in post-order. Let *g*_1_ and *g*_2_ denote the children of *g*. By Lemma 3.2, an rs-optimal reconciliation *R* between *G_g_* and *S* must extend some rs-optimal reconciliation *R*^1^ between *G*_*g*1_ and *S* and some rs-optimal reconciliation *R*^2^ between *G*_*g*2_ and *S*. Let **cost**(**ECrs**(·, ·, ·)) denote the cost component of an entry. Algorithm 1 describes how to complete table **ECrs**. Note that once all entries **ECrs**(·, ·, ·) have been computed, the optimal cost between *G* and *S* is simply 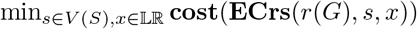.

#### Theorem 3.3.

*For each g* ∈ *V*(*G*), *s* ∈ *V*(*S*), *and* 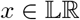, *Algorithm 1 correctly computes* **ECrs**(*g, s, x*).

#### Theorem 3.4.

*The time complexity of Algorithm 1 is O*(|*G*| · |*S*|^5^).

### 3.2 Reducing the Time Complexity

Next, we present an approach for speeding up the computation of table **ECrs** by a factor of O(|*S*|^3^). Our approach relies on the observation that many entries of **ECrs** will never correspond to an rs-optimal reconciliation and thus need not be considered in the dynamic program. Specifically, we show that for *g* ∈ *V*(*G*) and 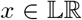, the set of species *s* ∈ *V*(*S*) that must be considered for entry **ECrs**(*g, s, x*) can be restricted to a set of constant size that corresponds to a generalization of the LCA.

Let *S_l_* denote the tree constructed from *S* with *e^L^* removed, and let *S_r_* denote the tree constructed from *S* with *e^R^* removed (Figure 2b). We extend the definition of the LCA to the *left lowest common ancestor*, denoted by *llca_S_*(*u, v*), and *right lowest common ancestor*, denoted by *rlca_S_*(*u, v*), defined as the lowest common ancestor of *u* and *v* in trees *S_l_* and *S_r_*. Let *BLCA_S_*(*u, v*) denote the set containing both the left and right lowest common ancestors.

Given a gene tree *G* and a species network *S* with one hybridization node, a reconciliation *R* between *G* and *S* is said to be a *BLCA mapping* if, for each internal node *g* of *G* with children *g*_1_ and *g*_2_, *R_v_*(*g*) ∈ *BLCA_S_*(*R_v_*(*g*_1_), *R_v_*(*g*_2_)). Note that if *G* has no internal nodes, then any reconciliation between *G* and *S* is trivially a BLCA mapping.

Let *R* and *R*\* be two reconciliations between *G* and *S*. *R*\* is said to *subsume R* if 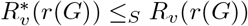, **signature**(*R*\*) ≤ **signature**(*R*), **edgeset**(*R*\*) ⊆ **edgeset**(*R*), and **DC**_*R*\*_ ≤ **DC**_*R*_ (Figure 2c).

#### Lemma 3.5.

*Given a gene tree G and a species network S with one hybridization node, let R* = (*R_v_, R_p_*) *be a reconciliation between G and S. If there exists an internal node g* ∈ *I*(*G*) *with children g*_1_ *and g*_2_ *such that R_v_*(*g*) ∉ *BLCA_S_*(*R_v_*(*g*_1_), *R_v_*(*g*_2_)), *then there exists some other reconciliation* 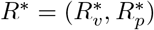 *between G and S such that for each u* ∈ *V*(*G*) *where g* ≤_*G*_ *u, R^*,u^ subsumes R^u^*.

#### Corollary 3.5.1.

*Given a gene tree G and species network S with one hybridization node, then for any reconciliation R* = (*R_v_, R_p_*) *between G and S that is not a BLCA mapping, there exists some other reconciliation R* that is a BLCA mapping and subsumes R*.

We are now ready to describe our revised dynamic programming algorithm for reconciling a gene tree *G* to a species network *S* with one hybridization node. In addition to **ECrs**, we construct a second table **candidates** that limits the set of species that need to be considered in completing **ECrs**. As in Algorithm 1, given any *g* ∈ *V*(*G*) and 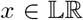, let *R* = (*R_v_, R_p_*) be an rs-optimal reconciliation between *G_g_* and *S* such that **signature**(*R*) = *x*. By Corollary 3.5.1, the algorithm need only consider *R* that are BLCA mappings; that is, for each internal node *g* with children *g*_1_ and *g*_2_, *R* must satisfy *R_v_*(*g*) ∈ *BLCA_S_*(*R_v_*(*g*_1_), *R_v_*(*g*_2_)). Let entry **candidates**(*g, x*) denote the set of possible values for *R_v_*(*G*), that is, the set of species nodes to which *g* can be mapped as part of some *R*. Then, for entry **ECrs**(*g, s, x*), it is clear that only entries for *s* ∈ **candidates**(*g, x*) need be computed. As before, entry **ECrs**(*g, s, x*) is an ordered pair (*E, c*), where *E* is the edgeset and *c* the cost of a reconciliation *R* as defined above. Algorithm 2 describes how to complete tables **candidates** and **ECrs**. Note that once all entries **candidates**(·, ·) and **ECrs**(·, ·) have been computed, the optimal cost between *G* and *S* is simply 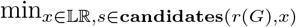 **cost**(**ECrs**(*r*(*G*), *s, x*)).

#### Theorem 3.6.

*For each g* ∈ *V*(*G*) *and* 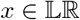, *Algorithm 2 correctly computes* **candidates**(*g, x*). *Furthermore, for each s* ∈ **candidates**(*g, x*), *Algorithm 2 correctly computes* **ECrs**(*g, s, x*).

#### Theorem 3.7.

*The time complexity of Algorithm 2 is* O(|*G*| · |*S*|^2^).

### 3.3 Allowing Gene Tree Handles

Next, we extend the previous results towards the ultimate goal of allowing for reconciliations with a level-1 species network. For reasons that will be clear later, for a gene tree *G*′ with root node *g*_1_ = *r*(*G*′), we consider the problem of reconciling a gene tree *G* with “handle” (*gh, g*_1_) to a species network *S* with one hybridization node. *gh* is a *handle node* that has no parents and a single child *g*_1_. Let *R* = (*R_v_, R_p_*) denote an rs-optimal reconciliation between *G* and *S* such that *R_v_*(*gh*) = *r*(*S*) and **signature**(*R*) = *x*. While *R* is not restricted to be a BLCA mapping, a reconciliation *R*′ between *G*′ and *S* is restricted to be a BLCA mapping. Algorithm 3 describes how to update **ECrs** accordingly via a straightforward modification of Algorithm 2.

#### Lemma 3.8.

*For each g* ∈ *V*(*G*) *and* 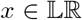, *Algorithm 3 correctly computes* **candidates**(*g, x*). *Furthermore, for each s* ∈ **candidates**(*g, x*), *Algorithm 3 correctly computes* **ECrs**(*g, s, x*).

#### Lemma 3.9.

*The time complexity of Algorithm 3 is* O(|*G*| · |*S*|^2^).

**Algorithm 2.**
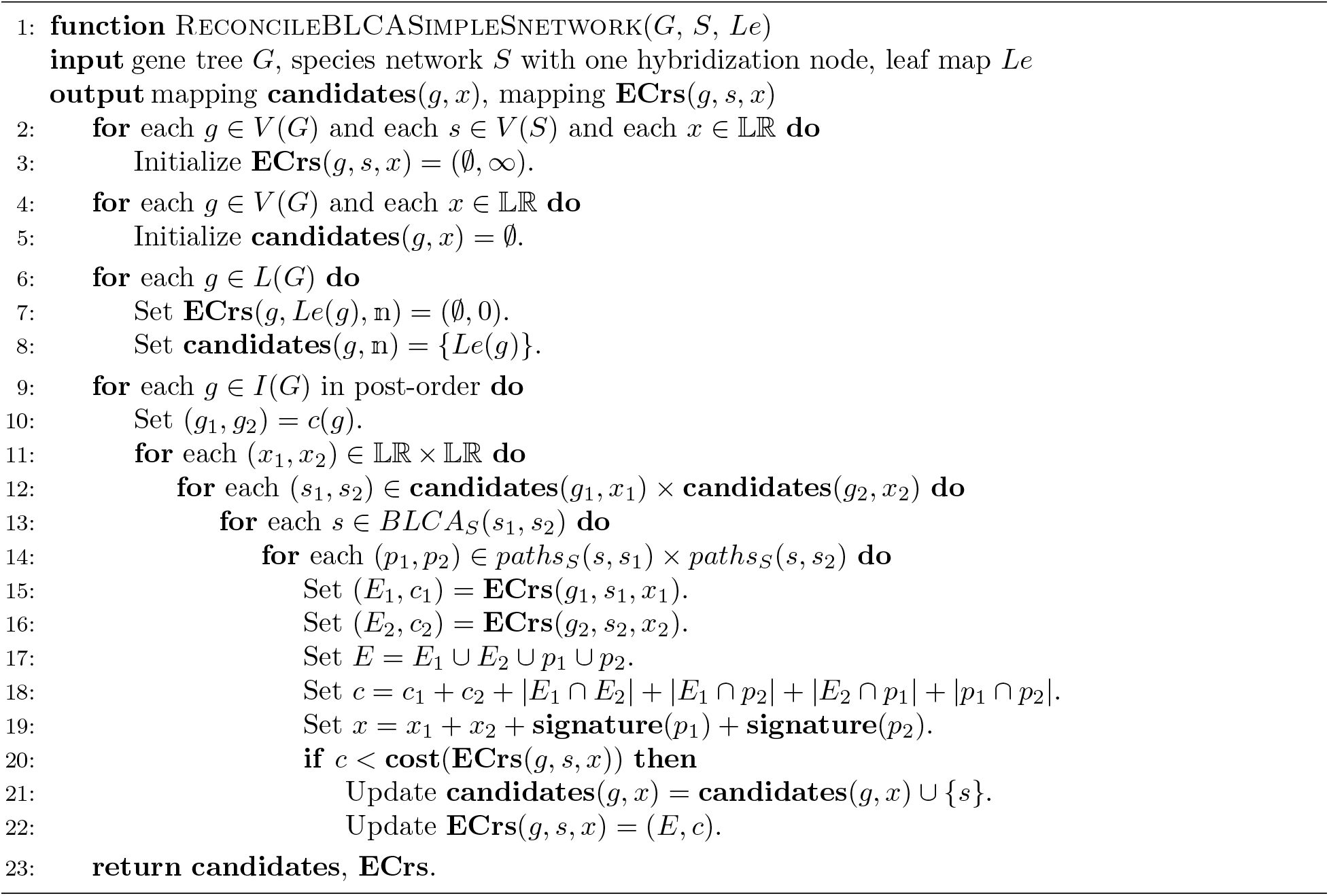

**Algorithm 3.**
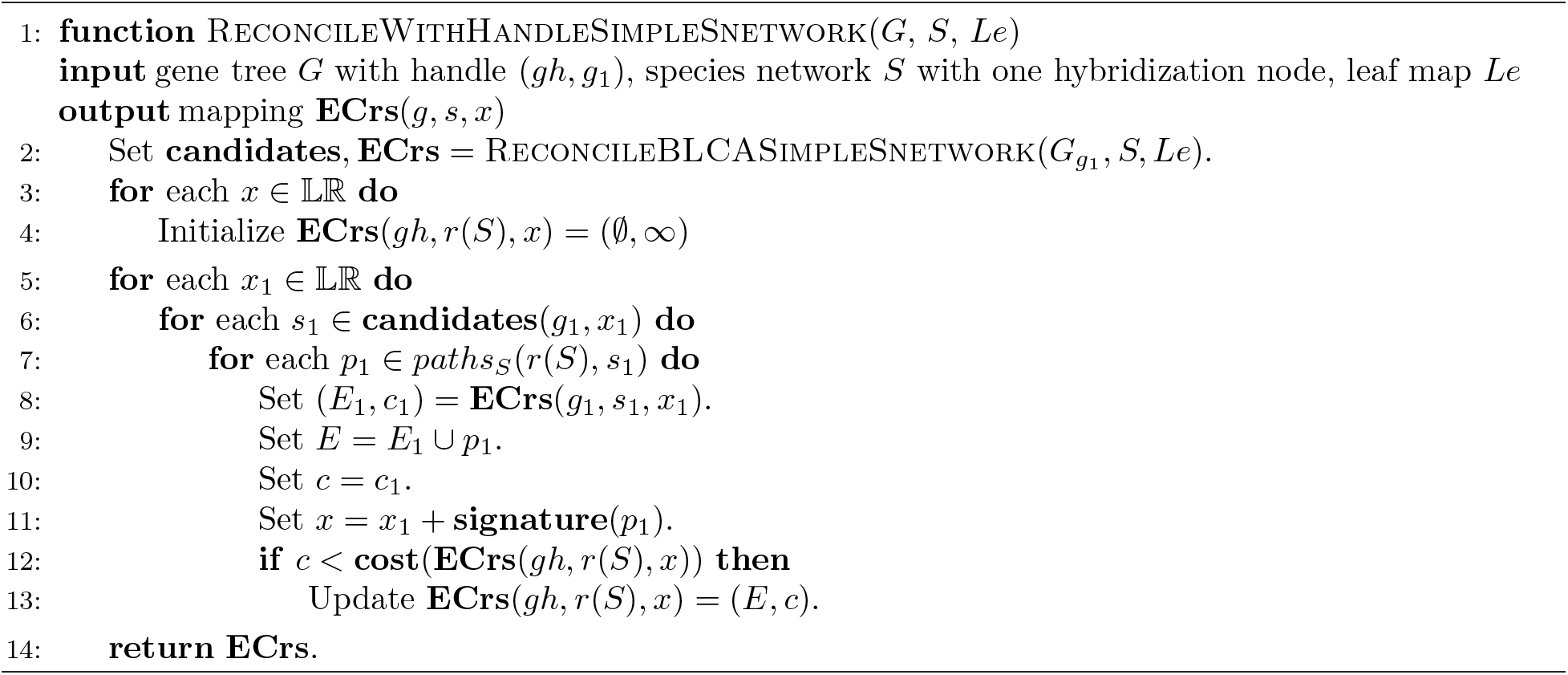

### 3.4 Considering Multiple Gene Trees

Next, we consider the problem of reconciling a forest of gene trees with handle nodes to a species network *S* with one hybridization node. We start by extending the definition of a reconciliation to a forest of gene trees.

#### Definition 3.1

(Forest Reconciliation). Let 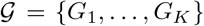 denote a forest of gene trees with handle nodes 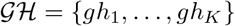. A *forest reconciliation* for 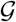 and *S* is a pair of mappings 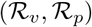 and a set of subreconciliations {*R*^1^,…, *R^K^*} subject to the following constraints:

1. For each *k* such that 1 ≤ *k* ≤ *K, R^k^* is a reconciliation between *G_k_* and *S*.
2. For each 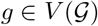, if *g* ∈ *V*(*G_k_*), then 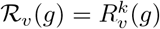 and 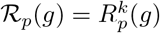.

Constraint 1 asserts that each *R^k^* is associated with *g_k_*, and Constraint 2 asserts that 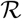 extends each *R^k^*.

For convenience, we often refer to a forest reconciliation as simply a reconciliation. To distinguish the two, we denote forest reconciliations using calligraphic font 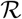 and (tree) reconciliations using standard math font *R*. In the remainder of this work, we include one additional constraint on all 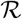: For each 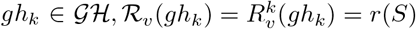. This constraint will be necessary later when we combine the forest of gene trees into a single tree. It is straightforward to extend definitions of edgeset, signature, lineages, and cost from (tree) reconciliations to forest reconciliations.

#### Lemma 3.10.

*Given a forest* 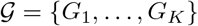 *of gene trees with handle nodes* 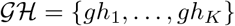 *and a species network S with one hybridization node, let* 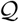 *and* 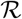 *denote two reconciliations between* 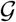 *and S. If* 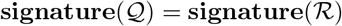, *then* 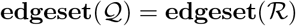.

Given a gene tree *G* with handle node *gh* and a species network *S* with one hybridization node, a reconciliation *R* = (*R_v_, R_p_*) between *G* and *S* is said to be *signature-optimal (s-optimal)* if it has the minimum cost among all reconciliations *R*′ between *G* and *S* such that 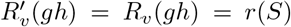 and **signature**(*R*′) = **signature**(*R*). Similarly, given a forest 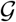 of gene trees with handle nodes and a species network *S*, a reconciliation 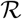 is said to be *signature-optimal (s-optimal)* if it has the minimum cost among all reconciliations 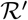 between 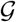 and *S* such that 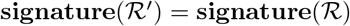.

Now, define some (arbitrary) order on the trees in a forest 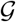. For each *k* such that 1 ≤ *k* ≤ *K*, let 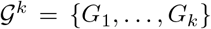 denote the first *k* gene trees of 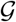 with handle nodes 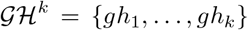. Let 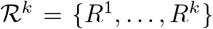 denote a reconciliation between 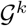 and *S*. It follows that 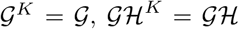, and 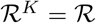.

#### Lemma 3.11

(Optimal Substructure Property for Forest of Trees). *Given a forest* 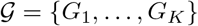 *of gene trees and a species network S with one hybridization node, let* 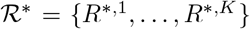 *denote an s-optimal reconciliation between* 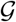 *and S. Then, for each k such that* 1 ≤ *k* ≤ *K*, 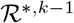 *and R^*,k^ are s-optimal*.

We are now ready to describe our dynamic programming algorithm for reconciling 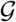 and *S*. Our algorithm constructs another table **ECs**. Given any 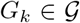 and 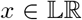, entry **ECs**(*G_k_, x*) is an ordered pair (*E, c*) for an s-optimal reconciliation 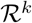 between 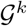 and *S* such that 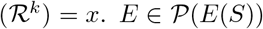 denotes the edgeset of 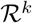, and non-negative integer *c* denotes the cost of 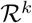. Note that by Lemma 3.10 and the definition of s-optimality, the specific reconciliation 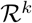 does not matter. As with **ECrs**, let **cost**(**ECs**(·, ·)) denote the cost component of an entry. Algorithm 4 describes how to complete table **ECs**. It is conceptually similar to Algorithm 1 but relies on Lemma 3.11 for multiple gene trees rather than Lemma 3.2 for a single gene tree. Once all entries have been computed, the algorithm returns the cost of reconciliation 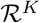.

#### Lemma 3.12.

*Algorithm 4 correctly computes the reconciliation cost*.

#### Lemma 3.13.

*The time complexity of Algorithm 4 is* 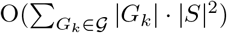.

**Algorithm 4.**
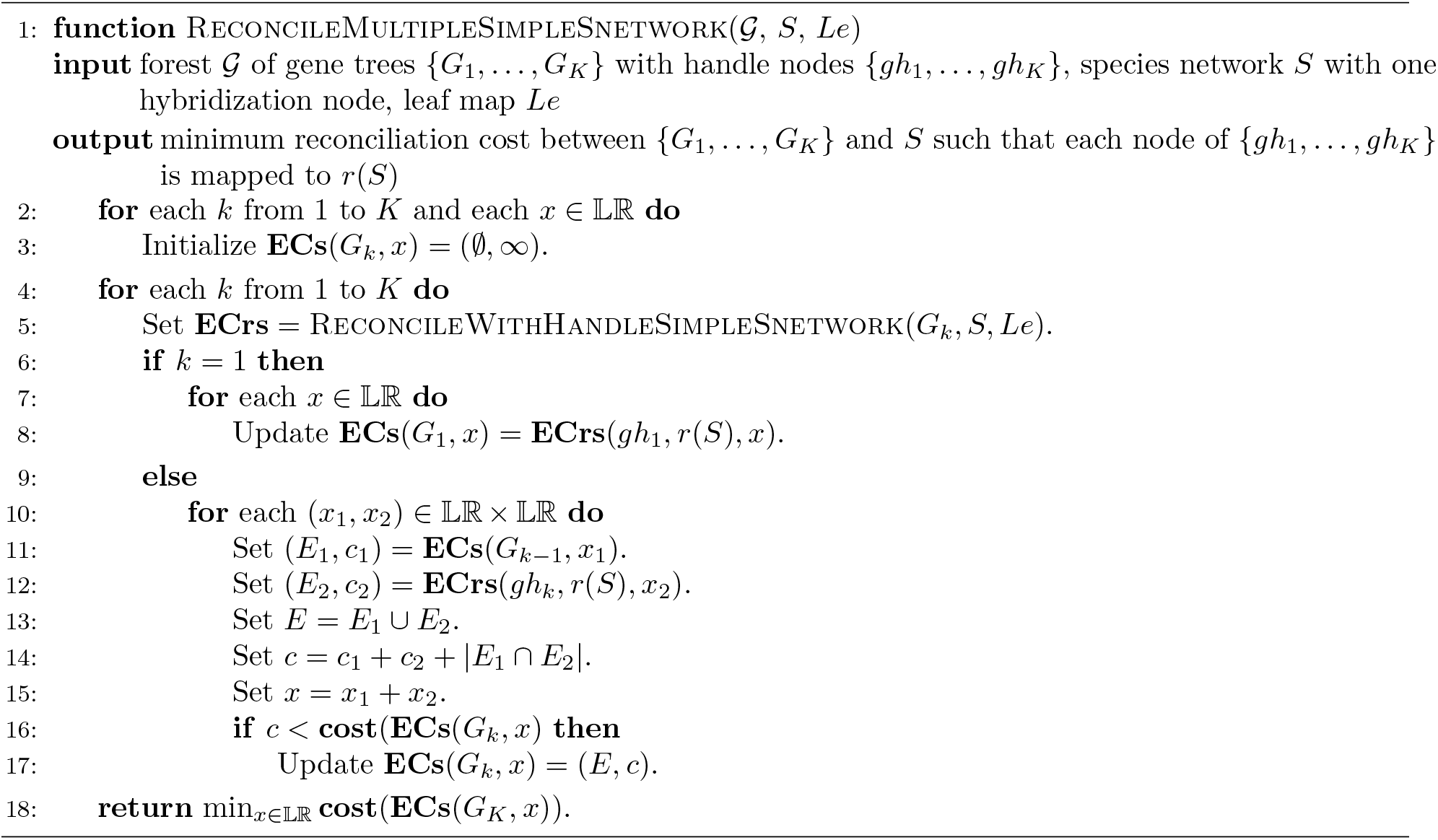

### 3.5 Putting the Pieces Together

In this section, we give an efficient algorithm for solving the most parsimonious reconciliation problem for a gene tree and a level-1 species network. This algorithm has some similarities with Algorithm 1 of To and Scornavacca (2015), which finds an optimal switching^3^ of a level-*k* species network that minimizes the duplication-loss cost between a gene tree and the resulting species tree. We demonstrate that their general approach of decomposing the gene tree and species network can be applied to our problem of minimizing the deep coalescence cost, where we reconcile each component of the decomposition using our previously presented algorithms.

We first give some definitions and lemmas, largely taken verbatim from To and Scornavacca (2015) except for minor modifications to notation.

Let *B* be a biconnected component of a network *S*. Then *B* contains exactly one node without ancestors in *B*; let *r*(*B*) denote the root of *B*. If *B* is not trivial, i.e. *B* consists of more than one node, we can contract it by removing all nodes of *B* other than *r*(*B*), then connect *r*(*B*) to every node with in-degree 0 created by this removal. Then the following definitions are well-posed (Figure 3).

**Figure 3.**
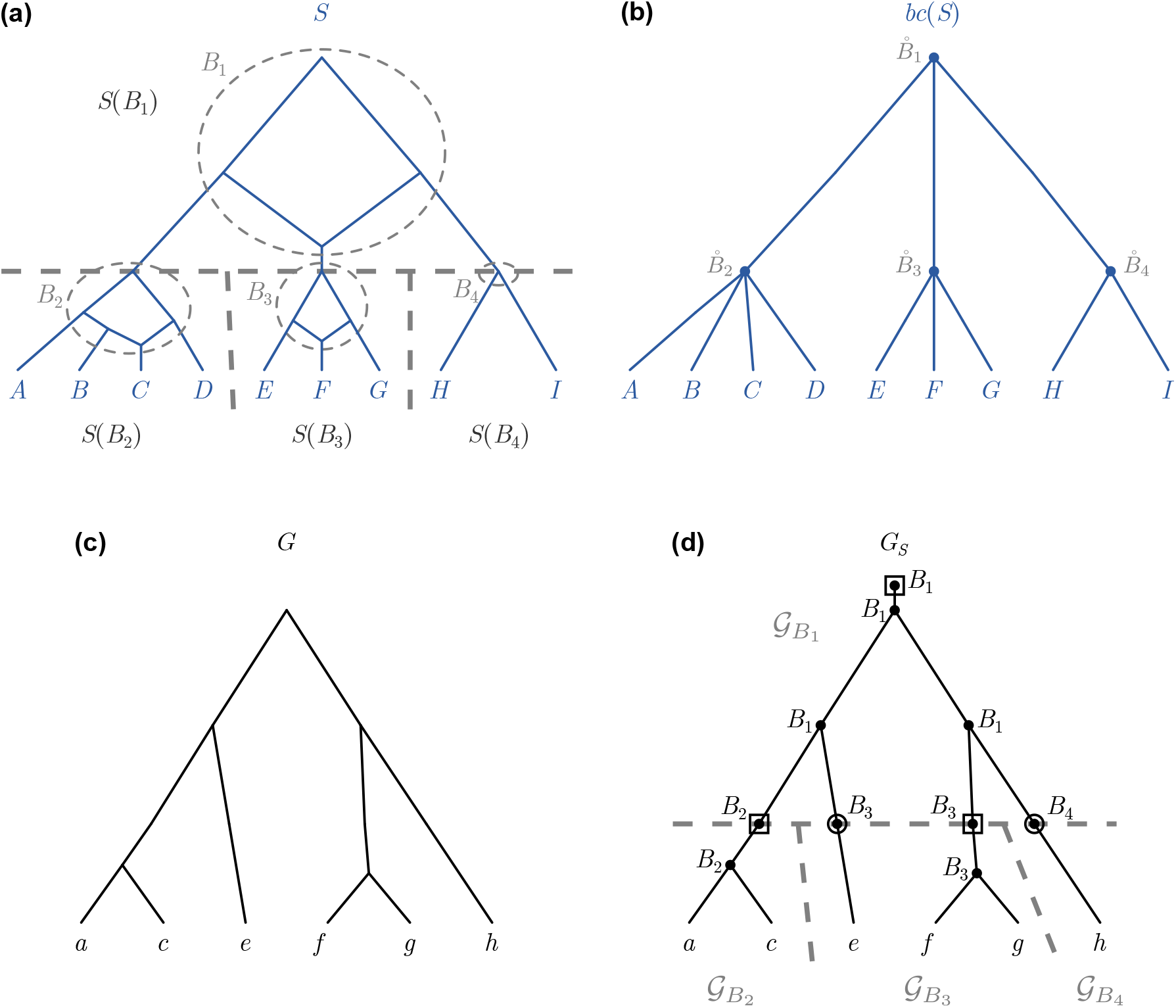
Annotated species networks and gene trees. (a) A level-1 species network *S* with four biconnected components *B_i_* and four elementary networks *S*(*B_i_*), where 1 ≤ *i* ≤ 4. (b) The tree *bc*(*S*) where, for each *i* such that 1 ≤ *i* ≤ 4, node 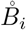 in *bc*(*S*) corresponds to biconnected component *B_i_* in *S*. (c) A gene tree *G*. (d) The tree *G_S_* along with its labeling 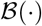 of nodes and subgraphs 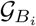, where 1 ≤ *i* ≤ 4. Circled nodes indicate artificial nodes (from Definition 3.5), and boxed nodes indicate handle nodes (from Definition 3.7). Figures and caption adapted from To and Scornavacca (2015).

#### Definition 3.2

(Tree *bc*(*S*); To and Scornavacca (2015), Definition 2). Given a network *S*, the tree *bc*(*S*) is obtained from *S* by contracting all its biconnected components.

Let 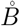 denote the node in *bc*(*S*) that corresponds to a biconnected component *B* in *S*. Given two biconnected component *B_i_* and *B_j_*, we say that *B_i_* ≤_*S*_ *B_j_* (resp. *B_i_* <_*S*_ *B_j_*) if and only if 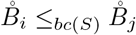 (resp. 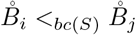). In such a case, *B_i_* is said to be *lower than or equal to* (resp. lower than) *B_j_*. We say that *B_i_* is the parent (resp. a child) of *B_j_* if 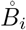 is the parent (resp. a child) of 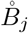 in *bc*(*S*).

#### Definition 3.3

(Elementary network; To and Scornavacca (2015), Definition 3). Given a network *S*, each biconnected component *B* that is not a leaf of *S* defines an elementary network, denoted by *S*(*B*), consisting of *B* and all edges (*u, v*) such that *u* ∈ *V*(*B*).^4^

#### Definition 3.4

(Mapping 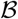; To and Scornavacca (2015), Definition 6). For every *u* ∈ *V*(*G*), 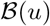 is defined as the lowest biconnected component *B* of *S* such that *L*(*S*_*r*(*B*)_) contains {*Le*(*v*) | *v* ∈ *L*(*G_u_*)}.^5^

#### Definition 3.5

(Tree *G_S_*; To and Scornavacca (2015), Definition 7). The tree *G_S_* is obtained from *G* as follows: For each internal node *u* in *G* with child nodes *u*_1_ and *u*_2_ such that there exist *k* biconnected components *B*_*i*_1__ >_*S*_… >_*S*_ *B_i_k__* strictly below 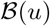 and strictly above 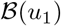, we add *k* artificial nodes *v*_1_ >… > *v_k_* on the edge (*u, u*_1_), and we fix 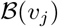 to *B_i_j__*. We do the same for *u*_2_.

#### Definition 3.6

(Subgraph 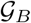; To and Scornavacca (2015), Definition 8). Let *B* be a biconnected component of *S* that is not a leaf. Then 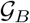 is the set of all maximal connected subgraphs *H* of *G_S_* such that 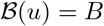 for every *u* ∈ *I*(*H*).

#### Lemma 3.14

(To and Scornavacca (2015), Lemma 2). *Let B be a biconnected component of S that is not a leaf. Then we have the following:*

i. *for every* 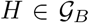, *H is either a binary tree or an edge whose upper extremity is an artificial node. Moreover, for every leaf u of H*, 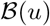 *is a child of B*.
ii. *if* 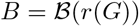, *then* 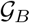 *consists of one binary tree*.

We make the following minor modifications to our above definitions and lemmas to require handles for each gene tree:

#### Definition 3.7

(Modified from Definitions 3.5 and 3.6). Obtain *G_S_* from *G* as in Definition 3.5. For each biconnected component *B* of *S* that is not a leaf, obtain 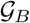 as in Definition 3.6. Then, consider each node *u* in *G* such that *u* is the root of some 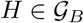 and *u* is not an artificial node. If 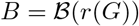, add a handle node *v* above *u*. Otherwise, add an artificial node *v* above *u*. Then, 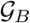 has a handle (*v, u*). Furthermore, fix 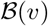 to 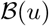.

#### Lemma 3.15

(Modified from Lemma 3.14). *Let B be as defined in Definition 3.14. Then we have the following:*

i. *for every* 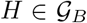, *H is either a binary tree **with a handle** or an edge whose upper extremity is an artificial node. Moreover, for every leaf u of H*, 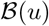 *is a child of B*.
ii. *if* 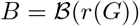, *then* 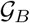 *consists of one binary tree **with a handle***.

It can be easily shown that adding artificial nodes and handle nodes to *G* does not change the minimum reconciliation cost between *G* and *S*.

Given a level-1 species networks *S*, a biconnected component of *S* is either a binary tree or a network with a single hybridization node. It is straightforward to modify each of our previous algorithms that reconcile one or more gene trees to a species network with one hybridization node, to reconcile one or more gene trees to a species tree (Supplemental Algorithms S1, S2, S3). The proofs of correctness and complexity are analogous to those of the corresponding algorithms and are therefore omitted.

Let 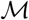 denote a mapping from nodes of *S* to biconnected components of *S* such that for each node *s* of *S*, 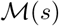 is the component to which *s* contracts. Given a gene tree *G*, a level-1 species network *S*, a mapping 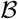, and a mapping 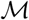, a reconciliation *R* between *G* and *S* is said to be *consistent* with 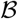 and 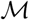 if, for each internal node *g* of *G* with children *g*_1_ and *g*_2_, 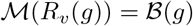. Note that if *G* has no internal nodes, then any reconciliation between *G* and *S* is trivially consistent.

#### Lemma 3.16.

*Given a gene tree G and a level-1 species network S, let R* = (*R_v_, R_p_*) *be a reconciliation between G and S. Given a mapping* 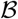 *and a mapping* 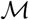, *if there exists an internal node g* ∈ *I*(*G*) *with children g*_1_ *and g*_2_ *such that* 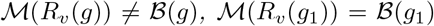, *and* 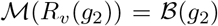, *then there exists some other reconciliation* 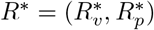 *between G and S such that R* subsumes R*.

#### Corollary 3.16.1.

*Given a gene tree G, a level-1 species network S, a mapping* 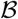, *and a mapping* 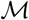, *then for any reconciliation R* = (*R_v_, R_p_*) *between G and S that is not consistent with* 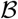 *and* 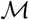, *there exists some other reconciliation R* that is consistent with* 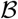 *and* 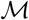 *and subsumes R*.

**Algorithm 5.**
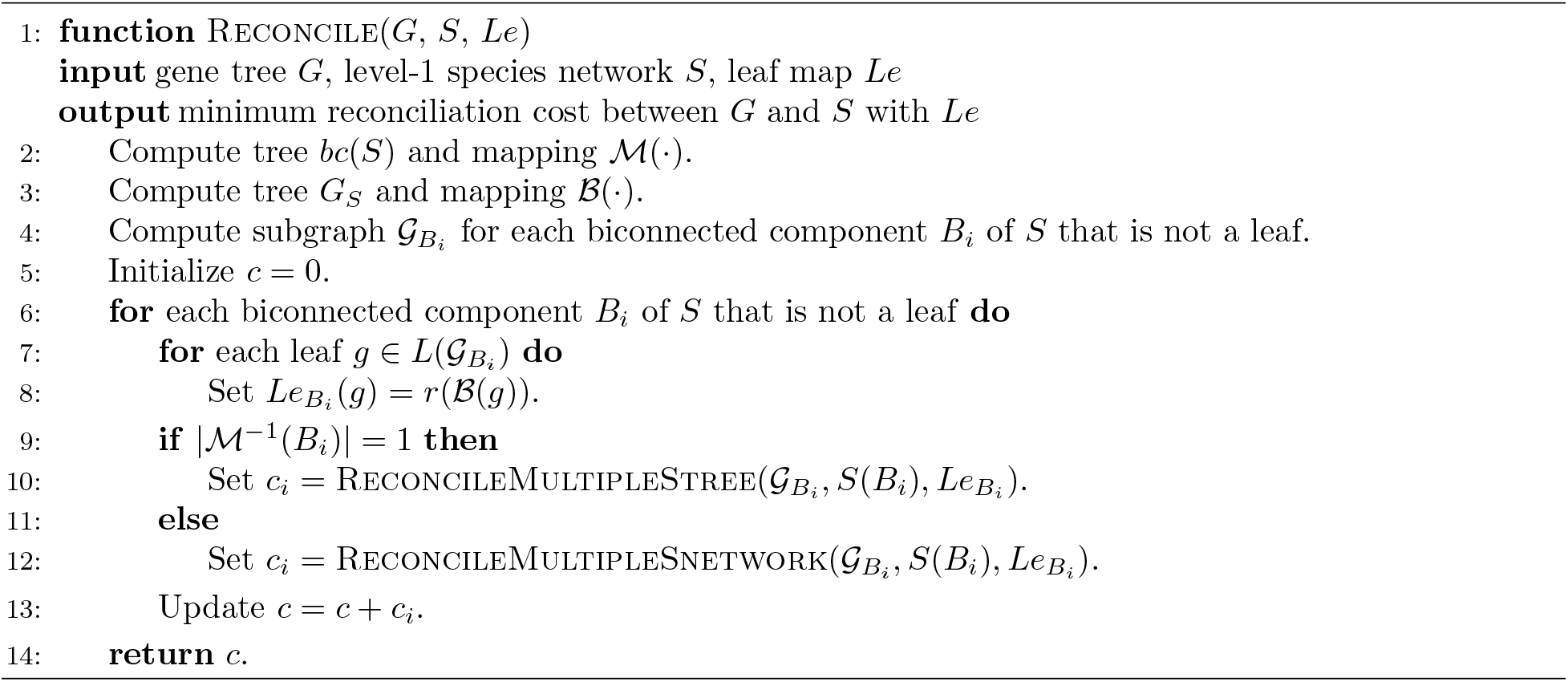

We are now ready to describe an algorithm for reconciling a binary gene tree *G* and a level-1 species network *S*. Algorithm 5 describes how to compute the minimum reconciliation cost between a *G* and *S* by analyzing each biconnected component of *S* independently (Figure 4).

**Figure 4.**
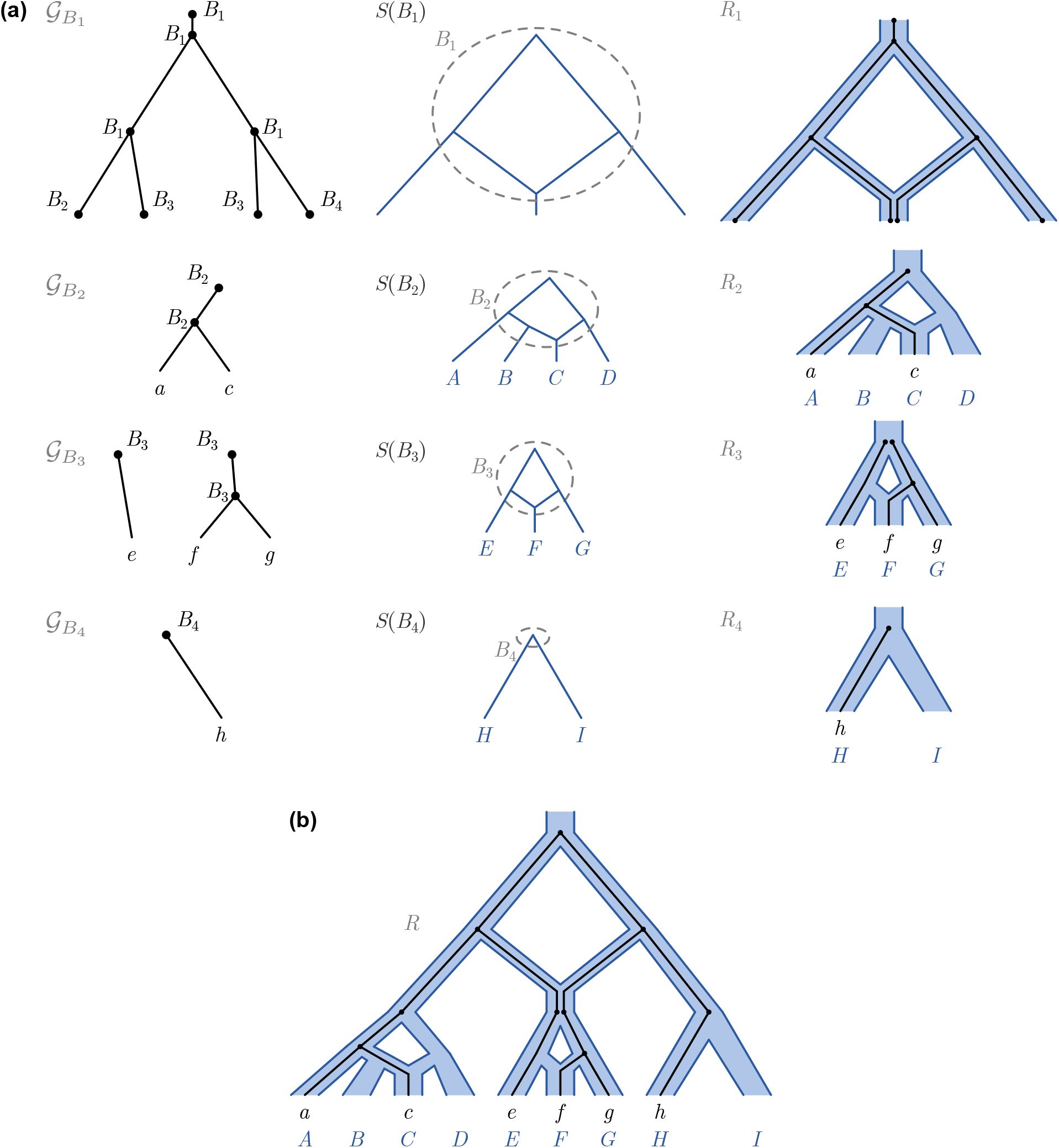
Reconciliation algorithm. (a) Continuing the example from Figure 3, reconciliations *R_i_* between subgraphs 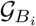 and *S*(*B_i_*), where 1 ≤ *i* ≤ 4. *R*_1_ induces 1 extra lineage, and *R*_2_, *R*_3_, and *R*_4_ each induce 0 extra lineages. (b) The full reconciliation between *G* and *S*.

#### Theorem 3.17.

*Algorithm 5 correctly computes the minimum reconciliation cost between G and S with leaf map Le*.

#### Theorem 3.18.

*The time complexity of Algorithm 5 is* O(|*G*| · |*S*|^2^).

### 3.6 Beyond Level-1 Networks

Algorithm 5 can be extended for general species networks *S* of level-*k*. To do so, Algorithms 2, 3, and 4 are easily extended to take species networks with up to *k* hybridization nodes. Such a modification requires tracking *k* separate signatures, one for each hybridization node. As there are four possible values for each signature, the time complexity of each extended algorithm would gain an additional factor of 4^*k*^, resulting in an overall time complexity of O(4^*k*^ · |*G*| · |*S*|^2^) for Algorithm 4. Since the complexity of Algorithm 4 dominates the complexity of Algorithm 5, the extended version of Algorithm 5 would then also have a time complexity of O(4^*k*^ · |*G*| · |*S*|^2^). Although this time complexity is exponential in the level of the network, we might expect *k* to be small for most phylogenies. Thus, the algorithm could still be practical in most cases.

## 4 Discussion

In this work, we have presented a polynomial-time algorithm for inferring most parsimonious reconciliations between gene trees and level-1 species networks that explain topological incongruence through hybridization and ILS. Our dynamic program required several developments. First, we introduced the concept of a reconciliation signature, which specifies which hybridization edges are used by different parts of the reconciliation. Next, we showed that the number of candidate species to consider in the dynamic program can be restricted to a set of constant size that corresponds to a generalization of the LCA. Finally, we decomposed the gene tree and species network using biconnected components and reconciled each component independently. While we have focused on level-1 networks, our algorithm can be extended to level-*k* species networks, though the time complexity is exponential in *k*.

We believe that our algorithm can be applied in several contexts. When the gene tree and species network are fixed, the algorithm can be used directly to infer reconciliations. Perhaps more interesting applications include incorporating the algorithm as part of a larger pipeline. For example, given a set of reconstructed gene trees, there exist several methods for species network inference using a parsimony criterion (Yu et al. 2011a; 2013a;b). However, since more complex networks (with more hybridization) can better fit the data (yielding reconciliations with equal or smaller numbers of extra lineages), methods are needed to balance this classic trade-off between complexity and fit. Lastly, if the species network is considered known but the gene alignments lack phylogenetic signal, reconciliation can be used to correct errors in gene tree topology (Wu et al. 2013, Bansal et al. 2015).

There are numerous directions for future work. There can exist multiple MPRs for a fixed gene tree and species network, and reconciliations are sensitive to user-defined event costs. While several papers have investigated the space of MPRs under the duplication-transfer-loss model (Bansal et al. 2013, Libeskind-Hadas et al. 2014) and the duplication-transfer-coalescence model (Du et al. 2019a, Mawhorter et al. 2019), we believe that similar problems can be explored in a joint hybridization and ILS model. Similarly, several reconciliation algorithms have been extended to handle non-binary gene trees (Yu et al. 2011b, Kordi and Bansal 2019) or species networks, or to incorporate macro-evolutionary events such as gene duplication, loss, and transfer (To and Scornavacca 2015, Du et al. 2019b). While hybridization and gene transfer may result in similar types of incongruence, more investigation is needed to see how we might disentangle the two signals. Finally, a theoretical analysis might address whether the MPR problem under a hybridization and ILS model is NP-hard for level-*k* species networks for arbitrary values of *k*.

## Acknowledgements

This work was supported by the Department of Computer Science and the Dean of Faculty of Harvey Mudd College. This material is based upon work supported by the National Science Foundation under Grant No. IIS-1751399 to YW and IIS-1905885 to RLH.

## Footnotes

1 When explaining topological incongruence through only deep coalescence, a reconciliation is sometimes called a *coalescent history* (Elworth et al. 2019).

2 Specifically, *R_v_* is the LCA reconciliation, and *R_p_* can be inferred from *R_v_*.

3 Per Definition 4 of To and Scornavacca (2015), a switching chooses, “for each hybridization edge, an incoming edge to switch on and the other to switch off.”

4 To and Scornavacca (2015) defines *S*(*B*) as “consisting of *B* and all cut-edges coming out from *B*”.

5 To and Scornavacca (2015) denoted this mapping as *B* and phrased the definition in terms of leaf labels. We use 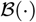 to distinguish the mapping from a biconnected component *B*.

## Notes

### Competing Interest Statement

The authors have declared no competing interest.

### Summary of Updates

Draft remarks removed.

