## Supplementary Material for "A Polynomial-Time Algorithm for Minimizing the Deep Coalescence Cost for Level-1 Species Networks"

#### S1 Notation

| **Notation** | **Description** |
|---|---|
| **Network** $N$ | |
| $V(N)$ | Nodes of $N$ (also called vertices) |
| $E(N)$ | Edges of $N$ (also called branches) |
| $L(N)$ | Leaves of $N$ |
| $I(N)$ | Internal nodes of $N$ |
| $r(N)$ | Root node of $N$ |
| $c(v)$ | Children nodes of node $v \in V(N)$ |
| $p(v)$ | Parent nodes of node $v \in V(N)$ (a single node or a set of two nodes) |
| $e(v)$ | Edge $(p(v), v)$ of node $v \in V(N)$ if $v$ has a single parent |
| $N_v$ | Subnetwork of $N$ rooted at $v \in V(N)$ |
| $\leq_N$ | Partial order on $V(N) \cup E(N)$ |
| $paths_N(v, u)$ | Set of paths from node $v \in V(N)$ to node $u \in V(N)$ in $N$ |
| $paths(N)$ | Set of all paths in $N$ |
| **Phylogenetic Reconciliation** | |
| $G$ | Gene tree |
| $S$ | Species network |
| $Le$ | Leaf map from $L(G)$ to $L(S)$ |
| $R$ | Reconciliation between $G$ and $S$ with leaf map $Le$ |
| $R_v$ | Vertex mapping of $R$ from $V(G)$ to $V(S)$ |
| $R_p$ | Path mapping of $R$ from $V(G)$ to $paths(S)$ |
| $R^g$ | Reconciliation $R$ restricted to subtree $G_g$ |
| $\mathbf{L}_R(e)$ | Number of lineages exiting edge $e \in E(S)$ in $R$ |
| $\mathbf{XL}_R(e)$ | Number of extra lineages exiting edge $e \in E(S)$ in $R$ |
| $\mathbf{DC}_R$ | Deep coalescence cost of $R$ |
| $\mathbf{edgeset}(R)$ | Set of species edges used in $R_p$ |
| $\mathcal{G}$ | Forest of gene trees |
| $\mathcal{R}$ | Forest reconciliation between $\mathcal{G}$ and $S$ with leaf map $Le$ |

**Table S1.1.** Summary of Basic Notations.

| Notation | Description |
|---|---|
| **Species Network with One Hybridization Node** | |
| $\mathbb{LR}$ | Set of four symbols $\{\mathtt{n}, \mathbb{l}, \mathtt{r}, \mathbb{b}\}$ that denote hybridization edges in a set |
| **signature**$(E)$ | Signature of a set $E \subseteq E(S)$ of edges |
| **signature**$(R)$ | Signature of a reconciliation $R$ |
| $v^H$ | Hybridization node |
| $v^L$ and $v^R$ | Left and right parents of hybridization node |
| $e^L$ and $e^R$ | Left and right edges to hybridization node |
| $v^A$ | Split node |
| $v^{AL}$ and $v^{AR}$ | Left and right children of split node |
| $S^O$ | Outside tree of $S$ (subgraph of $S$ that contains the set of nodes $u \in V(S)$ such that $u \not\leq_S v^A$, and all edges between these nodes) |
| $S^L$ | Left tree of $S$ (subgraph of $S$ that contains the set of nodes $u \in V(S)$ such that $u \leq_S v^{AL}$ and $u \not\leq_S v^L$, and all edges between these nodes) |
| $S^R$ | Right tree of $S$ (subgraph of $S$ that contains the set of nodes $u \in V(S)$ such that $u \leq_S v^{AR}$ and $u \not\leq_S v^R$, and all edges between these nodes) |
| $S^H$ | Hybrid tree of $S$ (subgraph of $S$ that contains the set of nodes $u \in V(S)$ such that $u \leq_S v^H$, and all edges between these nodes) |
| **Dynamic Programming Tables** | |
| **ECrs**$(g, s, x)$ | Ordered pair $(E, c)$ for an rs-optimal reconciliation $R = (R_v, R_p)$ between $G_g$ and $S$ such that $R_v(g) = s$ and **signature**$(R) = x$, where $E$ denotes the edgeset of $R$ and $c$ denotes the cost of $R$ |
| **candidates**$(g, x)$ | Set of species nodes to which $g$ can be mapped as part of some rs-optimal reconciliation $R$ between $G_g$ and $S$ such that **signature**$(R) = x$ and $R$ is a BLCA mapping |
| **ECs**$(G_k, x)$ | Ordered pair $(E, c)$ for an s-optimal reconciliation $\mathcal{R}^k$ between $\mathcal{G}^k$ and $S$ such that **signature**$(\mathcal{R}^k) = x$, where $E$ denotes the edgeset of $\mathcal{R}^k$ and $c$ denotes the cost of $\mathcal{R}^k$ |
| **Biconnected Components (adapted from To and Scornavacca (2015))** | |
| $bc(S)$ | Tree obtained from network $S$ by contracting all its biconnected components |
| $S(B)$ | Network consisting of biconnected component $B$ and all cut-edges coming out from $B$ |
| $\mathcal{B}$ | Mapping from nodes of $G$ to lowest biconnnected components of $S$ |
| $G_S$ | Tree obtained from $G$ with artificial nodes and handle nodes |
| $\mathcal{G}_B$ | Subgraph of $G$ consisting of all maximal connected subgraphs $H$ of $G_S$ such that $\mathcal{B}(u) = B$ for every $u \in I(H)$ |
| $\mathcal{M}$ | Mapping from nodes of $S$ to biconnected components of $S$ |

**Table S1.2.** Summary of Notations used in Algorithms.

### S2 Proofs

#### S2.1 Starting with a Simpler Problem

##### S2.1.1 Proof of Lemma 3.1

*Proof.* Let $s = R_v^1(r(G)) = R_v^2(r(G))$ and $x = \textbf{signature}(R^1) = \textbf{signature}(R^2)$. Let $\hat{L}$ denote the image of $L(G)$ under $Le$, that is, the set of leaves of $S$ that are mapped from the leaves of $G$.

**Case 1**: $x = \mathbb{b}$
In reconciliation $R^1$, consider the unique path from $r(G)$ to any leaf $g \in L(G)$ through the sequence of nodes $g_1 \geq_G \ldots \geq_G g_k$ where $g_1 = r(G)$ and $g_k = g$. Then the sequence of nodes $R_v^1(g_1) \geq_S \ldots \geq_S R_v^1(g_k)$ defines some path $p$ from $s$ to $v = R_v^1(g) \in \hat{L}$. Specifically, $p$ is equal to the joining of paths $R_p(g_2), \ldots, R_p(g_k)$. (Note that $R_p(g_1)$ is not included because it specifies the path to take in $S$ between $g_1$ and its parent.) It is easily shown that each edge $e$ on $p$ is in $\textbf{edgeset}(R^1)$, and that each $e$ in $\textbf{edgeset}(R^1)$ is part of some path $p$ from $s$ to $v \in \hat{L}$.

Since $S$ has a single hybridization node, there exists at most two distinct paths from $s$ to a node $v \in \hat{L}$. Since $x = \mathbb{b}$, $\textbf{edgeset}(R^1)$ is the union of the following sets:

$E_1$: edges in the unique path from $s$ to $v^A$,

$E_2$: edges in the unique path from $v^A$ to $v^L$ using edge $e^L$,

$E_3$: edges in the unique path from $v^A$ to $v^R$ using edge $e^R$,

$E_4$: edges in the unique paths from $v^A$ to each $v \in \hat{L}$ that is not a descendant of $v^H$, and

$E_5$: edges in the unique paths from $v^H$ to each $v \in \hat{L}$ that is a descendant of $v^H$.

By the same reasoning, $\textbf{edgeset}(R^2)$ comprises the same set of edges.

**Case 2**: $x \neq \mathbb{b}$
The claim follows as in the proof for $x = \mathbb{b}$, with the following modifications:

1. There exists a unique path from $s$ to a node $v \in \hat{L}$.
2. If $x = \mathbb{n}$, then $R^1$ can use neither $e^L$ nor $e^R$. Thus, $\textbf{edgeset}(R^1) = E_1 \cup E_4 \cup E_5$.
3. If $x = \mathbb{l}$, then $R^1$ must use $e^L$ at least once and cannot use $e^R$. Thus, $\textbf{edgeset}(R^1) = E_1 \cup E_2 \cup E_4 \cup E_5$.
4. If $x = \mathbb{r}$, then $R^1$ muse use $e^R$ at least once and cannot use $e^L$. Thus, $\textbf{edgeset}(R^1) = E_1 \cup E_3 \cup E_4 \cup E_5$.

Again, the same reasoning applies to $R^2$, and thus $\textbf{edgeset}(R^1) = \textbf{edgeset}(R^2)$. □

##### S2.1.2 Proof of Lemma 3.2

*Proof.* Suppose to the contrary that $R^*$ is an rs-optimal reconciliation such that $R^{*,g}$ is not rs-optimal. Then, let $R' = (R'_v, R'_p)$ denote an rs-optimal reconciliation between $G_g$ and $S$ such that $R'_v(g) = R_v^{*,g}(g)$ and $\textbf{signature}(R') = \textbf{signature}(R^{*,g})$. Consider a new reconciliation $R = (R_v, R_p)$ between $G$ and $S$ that is equal to $R'$ for the subtree $G_g$ and otherwise equal to $R^*$:

$$R_v(u) = \begin{cases} R'_v(u), & \text{if } u \in V(G_g) \\ R_v^*(u), & \text{otherwise} \end{cases} \qquad R_p(u) = \begin{cases} R'_p(u), & \text{if } u \in V(G_g) \\ R_p^*(u), & \text{otherwise} \end{cases}$$

Note that, by construction, $R'_v(g) = R_v^{*,g}(g) = R_v^*(g)$, and thus, $R$ is a valid reconciliation.

Next, define the following sets:

$$E_R^{in} = \bigcup_{u \in V(G_g)} R_p(u), \qquad E_R^{out} = \bigcup_{u \in V(G) \setminus V(G_g)} R_p(u),$$

and given an edge $e$ of $S$,

$$U_R^{in}(e) = \{u \in V(G_g) : e \in R_p(u)\}, \qquad U_R^{out}(e) = \{u \in V(G) \setminus V(G_g) : e \in R_p(u)\}.$$

That is, $E_R^{in}$ denotes the edgeset of $R$ restricted to $G_g$ (i.e. from $R'$), and $U_R^{in}(e)$ denotes the set of gene nodes such that $(p(u), u)$ is a lineage that exits $e$ for $R$ restricted to $G_g$. Similarly, $E_R^{out}$ and $U_R^{in}$ denote the corresponding sets for $R$ not restricted to $G_g$ (i.e. from $R^*$). Clearly, $U_R^{in}(e)$ and $U_R^{out}(e)$ are disjoint, $U_R^{in}(e) = 0$ for $e \notin E_R^{in}$, and $U_R^{out}(e) = 0$ for $e \notin E_R^{out}$. Let $\mathbf{L}_R^{in}(e) = |U_R^{in}(e)|$, and let $\mathbf{L}_R^{out}(e) = |U_R^{out}(e)|$.

Then the cost of $R$ is given by

$$\begin{aligned}
\mathbf{DC}_R &= \sum_{e \in \mathbf{edgeset}(R)} (\mathbf{L}_R(e) - 1) \\
&= \sum_{e \in E_R^{in} \setminus E_R^{out}} \left(\mathbf{L}_R^{in}(e) - 1\right) + \sum_{e \in E_R^{out} \setminus E_R^{in}} \left(\mathbf{L}_R^{out}(e) - 1\right) + \sum_{e \in E_R^{in} \cap E_R^{out}} \left(\mathbf{L}_R^{in}(e) + \mathbf{L}_R^{out}(e) - 1\right) \\
&= \left( \sum_{e \in E_R^{in}} \left(\mathbf{L}_R^{in}(e) - 1\right) - \sum_{e \in E_R^{in} \cap E_R^{out}} \left(\mathbf{L}_R^{in}(e) - 1\right) \right) \\
&\quad + \left( \sum_{e \in E_R^{out}} \left(\mathbf{L}_R^{out}(e) - 1\right) - \sum_{e \in E_R^{in} \cap E_R^{out}} \left(\mathbf{L}_R^{out}(e) - 1\right) \right) \\
&\quad + \left( \sum_{e \in E_R^{in} \cap E_R^{out}} \left(\mathbf{L}_R^{in}(e) - 1\right) + \sum_{e \in E_R^{in} \cap E_R^{out}} \left(\mathbf{L}_R^{out}(e) - 1\right) + \sum_{e \in E_R^{in} \cap E_R^{out}} 1 \right) \\
&= \sum_{e \in E_R^{in}} \left(\mathbf{L}_R^{in}(e) - 1\right) + \sum_{e \in E_R^{out}} \left(\mathbf{L}_R^{out}(e) - 1\right) + |E_R^{in} \cap E_R^{out}|.
\end{aligned}$$

Similarly, the cost of $R^*$ is given by

$$\mathbf{DC}_{R^*} = \sum_{e \in E_{R^*}^{in}} \left(\mathbf{L}_{R^*}^{in}(e) - 1\right) + \sum_{e \in E_{R^*}^{out}} \left(\mathbf{L}_{R^*}^{out}(e) - 1\right) + |E_{R^*}^{in} \cap E_{R^*}^{out}|.$$

Next, note that $E_R^{in} = \bigcup_{u \in V(G_g)} R_p(u) = \bigcup_{u \in V(G_g)} R'_p(u) = \mathbf{edgeset}(R')$, and similarly, $E_{R^*}^{in} = \bigcup_{u \in V(G_g)} R_p^*(u) = \mathbf{edgeset}(R^{*,g})$. Furthermore, by construction, $R'_v(g) = R_v^{*,g}(g)$ and $\mathbf{signature}(R') = \mathbf{signature}(R^{*,g})$, and thus, from Lemma 3.1, $\mathbf{edgeset}(R') = \mathbf{edgeset}(R^{*,g})$. Therefore, $E_R^{in} = \mathbf{edgeset}(R') = \mathbf{edgeset}(R^{*,g}) = E_{R^*}^{in}$. Since $R$ is equal to $R^*$ for $u \notin V(G_g)$, $E_R^{out} = E_{R^*}^{out}$, and for $e \in E_R^{out}$, $\mathbf{L}_R^{out}(e) = \mathbf{L}_{R^*}^{out}(e)$. Therefore,

$$\begin{aligned}
\mathbf{DC}_{R^*} - \mathbf{DC}_R &= \sum_{e \in E_{R^*}^{in}} \left(\mathbf{L}_{R^*}^{in}(e) - 1\right) - \sum_{e \in E_R^{in}} \left(\mathbf{L}_R^{in}(e) - 1\right) \\
&= \sum_{e \in \mathbf{edgeset}(R^{*,g})} \left(\mathbf{L}_{R^{*,g}}^{in}(e) - 1\right) - \sum_{e \in \mathbf{edgeset}(R')} \left(\mathbf{L}_{R'}^{in}(e) - 1\right) \\
&= \mathbf{DC}_{R^{*,g}} - \mathbf{DC}_{R'}\,.
\end{aligned}$$

By assumption, $R^{*,g}$ is not rs-optimal and $R'$ is rs-optimal, so it follows that $\mathbf{DC}_{R^{*,g}} > \mathbf{DC}_{R'}$. Thus, $\mathbf{DC}_{R^*} > \mathbf{DC}_R$, which contradicts the rs-optimality of $R^*$. □

##### S2.1.3 Proof of Theorem 3.3

*Proof.* Our proof is by structural induction on $g$.

**Base Case**

For each $g \in L(G)$, consider an rs-optimal reconciliation $R$ between $G_g$ and $S$. In accordance with Definition 2.1, $g$ must map to $Le(g)$, that is, $R_v(g) = Le(g)$. Furthermore, by definition, $r(G_g) = g$, and so $R_p(g) = \emptyset$. This reconciliation requires neither hybridization edge of the species network, so **signature**$(R) = \mathbb{n}$, and $R$ has an empty edgeset and a cost of 0. These entries are assigned correctly after the execution of the *for* loop in lines 4-5.

**Inductive Case**
Let $g \in I(G)$ and $(g_1, g_2) = c(g)$. Let us assume that the entries **ECrs**$(g_1, s_1, x_1)$ and **ECrs**$(g_2, s_2, x_2)$ are computed correctly for each $s_1 \in V(S)$, $s_2 \in V(S)$, $x_1 \in \mathbb{LR}$, and $x_2 \in \mathbb{LR}$. Based on this induction hypothesis, we will show that the entries **ECrs**$(g, s, x)$ for each $s \in V(S)$ and $x \in \mathbb{LR}$ are computed correctly as well.

Let $R^1 = (R_v^1, R_p^1)$ be an rs-optimal reconciliation between $G_{g_1}$ and $S$ such that $R_v^1(g_1) = s_1$ and **signature**$(R^1) = x_1$. Then $R^1$ has edgeset $E_1$ and cost $c_1$ given by **ECrs**$(g_1, s_1, x_1)$. Similarly, let $R^2 = (R_v^2, R_p^2)$ be an rs-optimal reconciliation between $G_{g_2}$ and $S$ with an analogous edgeset $E_2$ and cost $c_2$ given by **ECrs**$(g_2, s_2, x_2)$. Now consider a reconciliation $R = (R_v, R_p)$ between $G_g$ and $S$ subject to the following constraints:

(i) $R_v(g) = s$ and **signature**$(R) = x$ for some $s \in V(S)$ such that $s_1 \leq_S s$ and $s_2 \leq_S s$ and some $x \in \mathbb{LR}$. This constraint is simply for notational convenience.

(ii) $R_p(g) = \emptyset$. This constraint is required by Definition 2.1 since $r(G_g) = g$.

(iii) $R$ is consistent with $R^1$ for the subtree $G_{g_1}$ and consistent with $R^2$ for the subtree $G_{g_2}$ *except* for the path mappings $R_p^1(g_1)$ and $R_p^2(g_2)$. Specifically,

$$R_v(u) = \begin{cases} R_v^1(u), & \text{if } u \in V(G_{g_1}) \\ R_v^2(u), & \text{if } u \in V(G_{g_2}) \end{cases} \qquad R_p(u) = \begin{cases} R_p^1(u), & \text{if } u \in V(G_{g_1}) \setminus \{g_1\} \\ R_p^2(u), & \text{if } u \in V(G_{g_2}) \setminus \{g_2\} \end{cases}$$

This constraint is required by Lemma 3.2 and asserts that $R$ extends $R^1$ and $R^2$.

(iv) $R_p(g_1) = p_1$ and $R_p(g_2) = p_2$ for some path $p_1$ from $s$ to $s_1$ and some path $p_2$ from $s$ to $s_2$. This constraint maps the two edges $(g, g_1)$ and $(g, g_2)$ of $G_g$ to paths $p_1$ and $p_2$ in the species network.

It is easily verified that $R$ is a valid reconciliation. Let $E$ and $c$ denote the edgeset and cost of $R$. For $s \in V(S)$, we first show that the edgeset $E$, signature $x$, and cost $c$ of $R$ are computed correctly, then show that **ECrs**$(g, s, x)$ stores the edgeset and cost for some reconciliation that is rs-optimal.

**Edgeset of** $R$
The edgeset of $R$ is given by

$$\begin{aligned} E = \textbf{edgeset}(R) &= \bigcup_{u \in V(G_g)} R_p(u) \\ &= \left( \bigcup_{u \in V(G_{g_1}) \setminus \{g_1\}} R_p(u) \right) \cup \left( \bigcup_{u \in V(G_{g_2}) \setminus \{g_2\}} R_p(g) \right) \cup R_p(g_1) \cup R_p(g_2) \cup R_p(g). \end{aligned}$$

Note that, for the first term, from Constraint (iii), $\bigcup_{u \in V(G_{g_1}) \setminus \{g_1\}} R_p(u) = \bigcup_{u \in V(G_{g_1}) \setminus \{g_1\}} R_p^1(u) = \bigcup_{u \in V(G_{g_1})} R_p^1(u) =$ **edgeset**$(R^1) = E_1$. Similarly, for the second term, $\bigcup_{u \in V(G_{g_2})} R_p(u) = E_2$. The third and fourth terms are given by Constraint (iv), and the last term is given by Constraint (ii). Thus,

$$E = E_1 \cup E_2 \cup p_1 \cup p_2,$$

as computed in line 14.

**Signature of $R$**
Similarly, the signature of $R$ is given by

$$\begin{aligned} x &= \mathbf{signature}(R) = \mathbf{signature}(E) \\ &= \mathbf{signature}(E_1 \cup E_2 \cup p_1 \cup p_2) \\ &= \mathbf{signature}(E_1) + \mathbf{signature}(E_2) + \mathbf{signature}(p_1) + \mathbf{signature}(p_2) \\ &= \mathbf{signature}(R_1) + \mathbf{signature}(R_2) + \mathbf{signature}(p_1) + \mathbf{signature}(p_2) \\ &= x_1 + x_2 + \mathbf{signature}(p_1) + \mathbf{signature}(p_2), \end{aligned}$$

as computed in line 16.

**Cost of $R$**
Finally, consider the cost of $R$:

$$c = \mathbf{DC}_R = \sum_{e \in \mathbf{edgeset}(R)} (\mathbf{L}_R(e) - 1)$$

Note that for any edge $e \in E$, the following holds:

(1) If $e \in E_1$, then $R^1$ contributes $\mathbf{L}_{R^1}(e)$ lineages to $\mathbf{L}_R(e)$.

(2) If $e \in E_2$, then $R^2$ contributes $\mathbf{L}_{R^2}(e)$ lineages to $\mathbf{L}_R(e)$.

(3) If $e \in p_1$, then $p_1$ contributes 1 lineage to $\mathbf{L}_R(e)$.

(4) If $e \in p_2$, then $p_2$ contributes 1 lineages to $\mathbf{L}_R(e)$.

Next, construct a partition over $E$ as follows: Recall that $E_1 = \mathbf{edgeset}(R^1)$ and $R_v^1(g_1) = s_1$, so each edge of $E_1$ is lower than $s_1$. Furthermore, $p_1$ is defined to be a path that ends at $s_1$. Thus, $E_1$ and $p_1$ share no edges. By similar reasoning, $E_2$ and $p_2$ share no edges. Then it can be easily shown that $E = \mathbf{edgeset}(R)$ can be partitioned into the following sets:

(i) $E_1 \setminus (E_2 \cup p_2)$

(ii) $E_2 \setminus (E_1 \cup p_1)$

(iii) $p_1 \setminus (E_2 \cup p_2)$

(iv) $p_2 \setminus (E_1 \cup p_1)$

(v) $E_1 \cap E_2$

(vi) $E_1 \cap p_2$

(vii) $E_2 \cap p_1$

(viii) $p_1 \cap p_2$

Then, expanding the cost of $R$ over this partition, we get the following:

(i) $$\sum_{e \in E_1 \setminus (E_2 \cup p_2)} (\mathbf{L}_R(e) - 1) = \sum_{e \in E_1 \setminus (E_2 \cup p_2)} (\mathbf{L}_{R^1}(e) - 1)$$

(ii) $$\sum_{e \in E_2 \setminus (E_1 \cup p_1)} (\mathbf{L}_R(e) - 1) = \sum_{e \in E_2 \setminus (E_1 \cup p_1)} (\mathbf{L}_{R^2}(e) - 1)$$

(iii) $$\sum_{e \in p_1 \setminus (E_2 \cup p_2)} (\mathbf{L}_R(e) - 1) = \sum_{e \in p_1 \setminus (E_2 \cup p_2)} (1 - 1) = 0$$

(iv) $$\sum_{e \in p_2 \setminus (E_1 \cup p_1)} (\mathbf{L}_R(e) - 1) = \sum_{e \in p_2 \setminus (E_1 \cup p_1)} (1 - 1) = 0$$

(v) $$\begin{aligned} &\sum_{e \in E_1 \cap E_2} (\mathbf{L}_R(e) - 1) = \sum_{e \in E_1 \cap E_2} (\mathbf{L}_{R^1}(e) + \mathbf{L}_{R^2}(e) - 1) \\ &= \sum_{e \in E_1 \cap E_2} (\mathbf{L}_{R^1}(e) - 1) + \sum_{e \in E_1 \cap E_2} (\mathbf{L}_{R^2}(e) - 1) + |E_1 \cap E_2| \end{aligned}$$

(vi) $\sum_{e \in E_1 \cap p_2} (\mathbf{L}_R(e) - 1) = \sum_{e \in E_1 \cap p_2} (\mathbf{L}_{R^1}(e) + 1 - 1) = \sum_{e \in E_1 \cap p_2} (\mathbf{L}_{R^1}(e) - 1) + |E_1 \cap p_2|$

(vii) $\sum_{e \in E_2 \cap p_1} (\mathbf{L}_R(e) - 1) = \sum_{e \in E_2 \cap p_1} (\mathbf{L}_{R^2}(e) + 1 - 1) = \sum_{e \in E_2 \cap p_1} (\mathbf{L}_{R^2}(e) - 1) + |E_2 \cap p_1|$

(viii) $\sum_{e \in p_1 \cap p_2} (\mathbf{L}_R(e) - 1) = \sum_{e \in p_1 \cap p_2} (1 + 1 - 1) = |p_1 \cap p_2|$

Combining (i), the first term of (v), and the first term of (vi):

$$\begin{aligned}\sum_{e \in E_1 \setminus (E_2 \cup p_2)} (\mathbf{L}_{R^1}(e) - 1) + \sum_{e \in E_1 \cap E_2} (\mathbf{L}_{R^1}(e) - 1) + \sum_{e \in E_1 \cap p_2} (\mathbf{L}_{R^1}(e) - 1) &= \sum_{e \in E_1} (\mathbf{L}_{R^1}(e) - 1) \\ &= \mathbf{DC}_{R^1} = c_1\end{aligned}$$

Similarly, combining (ii), the second term of (v), and the first term of (vii):

$$\begin{aligned}\sum_{e \in E_2 \setminus (E_1 \cup p_1)} (\mathbf{L}_{R^2}(e) - 1) + \sum_{e \in E_1 \cap E_2} (\mathbf{L}_{R^2}(e) - 1) + \sum_{e \in E_2 \cap p_1} (\mathbf{L}_{R^2}(e) - 1) &= \sum_{e \in E_2} (\mathbf{L}_{R^2}(e) - 1) \\ &= \mathbf{DC}_{R^2} = c_2\end{aligned}$$

Thus, after some rearrangement,

$$c = c_1 + c_2 + |E_1 \cap E_2| + |E_1 \cap p_2| + |E_2 \cap p_1| + |p_1 \cap p_2|,$$

as computed in line 15.

**Update of $\mathbf{ECrs}(g, s, x)$**
Lastly, note the following:

(i) the *for* loop at line 8 searches over all possible signatures $x_1 \in \mathbb{LR}$ for $R^1$ and $x_2 \in \mathbb{LR}$ for $R^2$,

(ii) the *for* loop at line 9 searches over all possible species $s_1 \in V(S)$ for $R^1_v(g_1)$ and $s_2 \in V(S)$ for $R^2_v(g_2)$,

(iii) the *for* loop at line 10 searches over all possible species $s \in V(S)$ for $R_v(g)$ such that $s_1 \leq_S s$ and $s_2 \leq_S s$, and

(iv) the *for* loop at line 11 searches over all possible paths $p_1$ and $p_2$ for $R_p(g_1)$ and $R_p(g_2)$.

Thus, at the end of the *for* loop at lines 6-18, the algorithm will have evaluated the edgeset $E$, signature $x$, and cost $c$ for every reconciliation $R$ between $G_g$ and $S$ subject to the previously stated constraints. To update $\mathbf{ECrs}(g, s, x)$, what remains is to retain only the edgeset $E$ and cost $c$ for some reconciliation that is rs-optimal with respect to a specific root $s = R_v(g)$ and signature $x = \mathbf{signature}(R)$. This filter is executed in lines 17 and 18. □

###### S2.1.4 Proof of Theorem 3.4

*Proof.* Note that for two nodes $u$ and $v$ of $S$, line 11 requires computing $path_S(u, v)$. The algorithm for computing single-source shortest-paths in a DAG requires time $O(|S|)$ and is executed $O(|S|)$ times, for a total complexity of $O(|S|^2)$.

Next, consider the time complexity of Algorithm 1 step-by-step. The *for* loop at lines 2-3 requires time $O(|G| \cdot |S|)$. The *for* loop at lines 4-5 requires time $O(|G|)$. The *for* loop at lines 6-18 is performed $O(|G|)$ times. At each iteration, each of $x_1$ and $x_2$ take on four values (line 8); each of $s_1$, $s_2$, and $s$ take on at most $|S|$ values (lines 9-10); and each of $p_1$ and $p_2$ take on at most two values (line 11). Thus, each of lines 12-18 is performed $O(|G| \cdot |S|^3)$ times. Since the number of edges in either edgeset $E$ or in path $p$ is bounded by $O(|S|)$, lines 14 and 15 each require at most time $O(|S|^2)$. Each other statement in lines 7-18 requires time $O(1)$. Therefore, lines 6-18 require time $O(|G| \cdot |S|^5)$, which subsumes the preprocessing time and the time for the other *for* loops. □

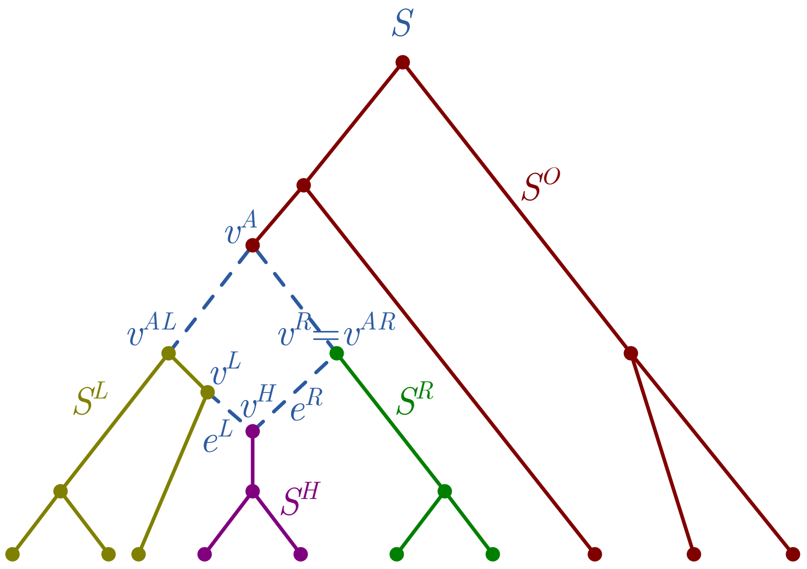


**Figure S2.1. Partition of species network with one hybridization node.**

#### S2.2 Reducing the Time Complexity

Let $v^H$, $v^L$, $v^R$, $e^L$, $e^R$, and $v^A$ be defined as before. Let $v^{AL}$ and $v^{AR}$ be the left and right children of $v^A$. There exists a partition of the nodes and edges of $S$ based on their position relative to $v^A$, $v^{AL}$, $v^{AR}$, and $v^H$ (Figure S2.1):

- the *outside set* contains the set of nodes $u \in V(S)$ such that $u \not\leq_S v^A$, and all edges between these nodes,
- the *left set* contains the set of nodes $u \in V(S)$ such that $u \leq_S v^{AL}$ and $u \not\leq_S v^L$, and all edges between these nodes,
- the *right set* contains the set of nodes $u \in V(S)$ such that $u \leq_S v^{AR}$ and $u \not\leq_S v^R$, and all edges between these nodes,
- the *hybrid set* contains the set of nodes $u \in V(S)$ such that $u \leq_S v^H$, and all edges between these nodes,
- the edge $(v^A, v^{AL})$ connects the outside set to the left set,
- the edge $(v^A, v^{AR})$ connects the outside set to the right set,
- the edge $e^L$ connects the left set to the hybrid set, and
- the edge $e^R$ connects the right set to the hybrid set.

Let $S^O$, $S^L$, $S^R$, and $S^H$ denote the subgraphs of $S$ comprising the outside, left, right, and hybrid sets, respectively, and let $\mathbb{S} = \{S^O, S^L, S^R, S^H\}$. It is easily verified that each $\hat{S} \in \mathbb{S}$ is a tree in $S$.

Here, we list several useful facts about the four trees comprising $\mathbb{S}$ and the various versions of lowest common ancestors, which can be easily verified:

(1) If $a$ and $b$ are in the same tree $\hat{S} \in \mathbb{S}$, then $lca_S(a,b) = llca_S(a,b) = rlca_S(a,b) \in V(\hat{S})$.

(2) If $a \in V(S^O)$ and $b \notin V(S^O)$, then $lca_S(a,b) = lca_S(a,v^A) = llca_S(a,b) = llca_S(a,v^A) = rlca_S(a,b) = rlca_S(a,v^A)$. In particular, for the case where $a = v^A$, it follows that $lca_S(b,v^A) = llca_S(b,v^A) = rlca_S(b,v^A) = v^A$.

(3) If $a \in V(S^L)$ and $b \in V(S^R)$, then $lca_S(a,b) = llca_S(a,b) = rlca_S(a,b) = v^A$

(4a) If $a \in V(S^L)$ and $b \in V(S^H)$, then $lca_S(a,b) = llca_S(a,b) \in V(S^L)$ and $rlca_S(a,b) = v^A$.

(4b) If $a \in V(S^R)$ and $b \in V(S^H)$, then $lca_S(a,b) = rlca_S(a,b) \in V(S^R)$ and $llca_S(a,b) = v^A$.

Given two paths $p_1$ and $p_2$ in $S$ with a common start node $s$, a *shared node* is defined to be a node common to both paths. The *lowest shared node* of $p_1$ and $p_2$, denoted $lsn_S(p_1, p_2)$, is defined to be the shared node $v$ such that there exists no shared node $u <_S v$.

**Lemma S2.1.** *Given a species network $S$ with one hybridization node, let $u$ and $v$ denote two nodes of $S$, and let $w$ denote a node of $S$ that is a common ancestor of $u$ and $v$. Then given a path $p_1$ from $w$ to $u$ and a path $p_2$ from $w$ to $v$, $lsn_S(p_1, p_2) \in BLCA_S(u, v)$.*

**Lemma S2.2.** *In Algorithm 2, each set* **candidates**$(g, x)$ *contains at most two elements.*

##### S2.2.1 Proof of Lemma S2.1

*Proof.* Given a tree $S'$ in $S$ and a path $p$ in $S$, $S'$ is said to *contain* $p$ if for each edge $e \in p$, $e \in E(S')$. Assume there exists a tree $S'$ in $S$ that contains both $p_1$ and $p_2$. Then $lsn_S(p_1, p_2) = lsn_{S'}(p_1, p_2)$, and $p_1$ and $p_2$ are the unique paths from $w$ to $u$ and $w$ to $v$, respectively, in $S'$. Let $X$ denote the set of shared nodes of $p_1$ and $p_2$. By definition, $X$ is also the set of common ancestors of $u$ and $v$ that are descendants of $w$. Thus, the lowest node in $X$ with respect to $\leq_{S'}$ is equal to $lsn_{S'}(p_1, p_2) = lca_{S'}(u, v)$.

Therefore, if $S_l$ contains both $p_1$ and $p_2$, then $lsn_S(p_1, p_2) = lca_{S_l}(u, v) = llca_S(u, v)$. And similarly, if $S_r$ contains both $p_1$ and $p_2$, then $lsn_S(p_1, p_2) = lca_{S_r}(u, v) = rlca_S(u, v)$.

Finally, if neither $S_l$ nor $S_r$ contain both $p_1$ and $p_2$, then the union of $p_1$ and $p_2$ includes both $e^L$ and $e^R$. Because both $e^L$ and $e^R$ lead to $v^H$, one of $p_1$ or $p_2$ contains $e^L$, the other contains $e^R$, and $v^H$ is a shared node of $p_1$ and $p_2$. Let $\hat{p}_1$ denote the subpath of $p_1$ from $v^H$ to $u$, and let $\hat{p}_2$ denote the subpath of $p_2$ from $v^H$ to $v$. Clearly, $lsn_S(\hat{p}_1, \hat{p}_2) = lsn_S(p_1, p_2)$. Next, note that $S^H$ contains both $\hat{p}_1$ and $\hat{p}_2$. Therefore, as before, $lsn_S(\hat{p}_1, \hat{p}_2) = lca_{S^H}(u, v)$, and since $S^H$ is a subtree of $S$, $lca_{S^H}(u, v) = lca_S(u, v)$. Finally, since both $u$ and $v$ are in $S^H$, by fact 1, $lca_S(u, v) = llca_S(u, v) = rlca_S(u, v)$. □

##### S2.2.2 Proof of Lemma 3.5

*Proof.* Let $s = R_v(g)$, $s_1 = R_v(g_1)$, and $s_2 = R_v(g_2)$. Let $p = R_p(g)$, $p_1 = R_p(g_1)$, and $p_2 = R_p(g_1)$. By assumption, $s \notin BLCA_S(s_1, s_2)$. By Definition 2.1, $p_1$ is a path from $s$ to $s_1$, and $p_2$ is a path from from $s$ to $s_2$. Let $d = lsn_S(p_1, p_2) \leq_S s$. By Lemma S2.1, $d \in BLCA_S(s_1, s_2)$.

Consider the following reconciliation $R^* = (R^*_v, R^*_p)$:

$$R^*_v(u) = \begin{cases} d, & \text{if } u = g \\ R_v(u), & \text{otherwise} \end{cases}$$

$$R^*_p(u) = \begin{cases} \emptyset & \text{if } u = g \text{ and } g = r(G) \\ p \text{ extended with the edges of } p_1 \text{ from } s \text{ to } d, & \text{if } u = g \text{ and } g \neq r(G) \\ p_1 \text{ truncated to start at } d, & \text{if } u = g_1 \\ p_2 \text{ truncated to start at } d, & \text{if } u = g_2 \\ R_p(u), & \text{otherwise} \end{cases}$$

Note that $R^*$ is a valid reconciliation.

Next, we consider for each $u \in V(G)$ such that $g \leq_G u$, the reconciliations $R^*$ and $R$ restricted to $G_u$. Specifically, we show that $R^{*,u}$ subsumes $R^u$. Note that since subtree $G_u$ does not include edge $e(u)$, for the restricted reconciliations, $R^{*,u}_p(u) = \emptyset$ and $R^u_p(u) = \emptyset$. Now compare $R^{*,u}$ and $R^u$. If $u = g$, then $R^{*,u}_v(u) = d \leq_S s = R^u_v(u)$, and otherwise, $R^{*,u}_v(u) = R^u_v(u)$. In creating $R^{*,u}_p$ from $R^u_p$, edges were either removed from paths or moved to different paths. Thus, **signature**$(R^{*,u}) \leq$ **signature**$(R^u)$, **edgeset**$(R^{*,u}) \subseteq$ **edgeset**$(R^u)$, and for each edge $e$ of $S$, $\mathbf{L}_{R^{*,u}}(e) \leq \mathbf{L}_{R^u}(e)$, so $\mathbf{DC}_{R^{*,u}} \leq \mathbf{DC}_{R^u}$. □

##### S2.2.3 Proof of Theorem 3.6

*Proof.* It suffices to show that for each $g \in V(G)$ and $x \in \mathbb{LR}$, the value of **candidates**$(g, x)$ is computed correctly. Then for each $s \in$ **candidates**$(g, x)$, the updates of **ECrs**$(g, s, x)$ have not changed from Algorithm 1, so the value of **ECrs**$(g, s, x)$ is computed correctly. Furthermore, **ECrs** contains only entries **ECrs**$(g, s, x)$ for $s \in$ **candidates**$(g, x)$, as required. Our proof is by structural induction on $g$.

**Base Case**
For each $g \in L(G)$, consider an rs-optimal reconciliation $R$ between $G_g$ and $S$. As in Theorem 3.3, it must be that $R_v(g) = Le(g)$ and **signature**$(R) = \mathbb{n}$. Thus, **candidates**$(g, \mathbb{n}) = \{Le(g)\}$, as computed in line 8.

**Inductive Case**
Let $g \in I(G)$ and $(g_1, g_2) = c(g)$. Let us assume that the entries **candidates**$(g_1, x_1)$ and **candidates**$(g_2, x_2)$ for each $x_1 \in \mathbb{LR}$ and $x_2 \in \mathbb{LR}$ are computed correctly. Furthermore, let us assume that the entries **ECrs**$(g_1, s_1, x_1)$ and **ECrs**$(g_2, s_2, x_2)$ for each $s_1 \in$ **candidates**$(g_1, x_1)$ and $s_2 \in$ **candidates**$(g_2, x_2)$ are computed correctly. Based on this induction hypothesis, we will show that the entries **candidates**$(g, x)$ for each $x \in \mathbb{LR}$ are computed correctly as well.

As in Theorem 3.3, let $R^1 = (R^1_v, R^1_p)$ be an rs-optimal reconciliation between $G_{g_1}$ and $S$ such that $R^1_v(g_1) = s_1$ and **signature**$(R^1) = x_1$, with edgeset $E_1$ and cost $c_1$ given by **ECrs**$(g_1, s_1, x_1)$. Similarly, let $R^2 = (R^2_v, R^2_p)$ be an rs-optimal reconciliation between $G_{g_2}$ and $S$ with an analogous edgeset $E_2$ and cost $c_2$ given by **ECrs**$(g_2, x_2)$. The inductive hypothesis guarantees $C(g_1, R^1_v(g_1), x_1)$ and $C(g_2, R^2_v(g_2), x_2)$ are computed correctly, since $R^1$ and $R^2$ must be BLCA mappings. Now consider an rs-optimal reconciliation $R = (R_v, R_p)$ between $G_g$ and $S$ subject to the same constraints as Theorem 3.3. By Corollary 3.5.1, it suffices to restrict $R$ to be a a BLCA mapping. That is, to update **candidates**$(g, x)$, it suffices to search over $s_1 \in$ **candidates**$(g_1, x_1)$, $s_2 \in$ **candidates**$(g_2, x_2)$, and $s \in BLCA_S(s_1, s_2)$, as in lines 12-13.

At the end of the *for* loop at lines 9-22, the algorithm will have evaluated the edgeset $E$, signature $x$, and cost $c$ of every rs-optimal reconciliation $R$ between $G_g$ and $S$ that is also a BLCA mapping. To update **candidates**$(g, x)$, what remains is to retain only the species $s$ such that there exists a reconciliation with $R_v(g) = s$ that is rs-optimal with respect to a specific root $s$ and signature $x =$ **signature**$(R)$. This filter is executed in lines 20- 21. □

##### S2.2.4 Proof of Lemma S2.2

*Proof.* We will prove a slightly stronger result by structural induction on $g$: Each set **candidates**$(g, x)$ is empty, contains one element, or contains two elements, one of which is $v^A$ and the other of which is in $S^L$ or $S^R$. Note that for $g \in V(G)$, if there exists no rs-optimal BLCA reconciliation $R$ between $G_g$ and $S$ such that **signature**$(R) = x$, then **candidates**$(g, x) = \emptyset$ as initialized (line 5).

**Base Case**
For $g \in L(G)$, **candidates**$(g, \mathbb{n})$ contains one element $Le(g)$ (line 8).

**Inductive Case**
Let $g \in I(G)$ and $(g_1, g_2) = c(g)$. Note that each $s$ added to an set **candidates**$(g, \cdot)$ (line 21) satisfies $s \in BLCA_S(s_1, s_2)$ (line 13), where $s_1 \in$ **candidates**$(g_1, x_1)$ and $s_2 \in$ **candidates**$(g_2, \cdot)$ (line 12). Assume by way of induction that each of **candidates**$(g_1, \cdot)$ and **candidates**$(g_2, \cdot)$ contains either one element or two elements, one of which is $v^A$ and the other of which is in $S^L$ or $S^R$. We consider three cases:

**Case 1**: **candidates**$(g_1, \cdot)$ and **candidates**$(g_2, \cdot)$.
Let $u_1$ denote the sole element of **candidates**$(g_1, \cdot)$, and let $u_2$ denote the sole element of **candidates**$(g_2, \cdot)$. Then **candidates**$(g, \cdot) = BLCA_S(u_1, u_2)$. Let $w = lca_S(u_1, u_2)$, $x = llca_S(u_1, u_2)$, and $y = rlca_S(u_1, u_2)$. Now consider the possible membership of $u_1$ and $u_2$ among the nodes of $S$.

If both $u_1$ and $u_2$ are nodes of the same tree $\hat{S} \in \mathbb{S}$, then from fact 1, $w = x = y$. Thus, $BLCA_S(u_1, u_2) = \{w\}$.

If one of $u_1$ or $u_2$ is a node of $S^O$ and the other is not, then by fact 2, $w = x = y$. Thus, $BLCA_S(u_1, u_2) = \{w\}$.

If one of $u_1$ or $u_2$ is a node of $S^L$ and the other is a node of $S^R$, then from fact 3, $w = x = y$. Thus, $BLCA_S(u_1, u_2) = \{w\}$.

If one of $u_1$ or $u_2$ is a node of $S^L$ and the other is a node of $S^H$, then from fact 4a, $w = x \in V(S^L)$ and $y = v^A$. Thus, $BLCA_S(u_1, u_2) = \{w, v^A\}$.

If one of $u_1$ or $u_2$ is a node of $S^R$ and the other is a node of $S^H$, then from fact 4b, $w = y \in V(S^R)$ and $x = v^A$. Thus, $BLCA_S(u_1, u_2) = \{w, v^A\}$.

In conclusion, either **candidates**$(g, \cdot) = \{w\}$, or **candidates**$(g, \cdot) = \{v^A, w\}$, where $w \in V(S^L) \cup V(S^R)$.

**Case 2**: **candidates**$(g_1, \cdot)$ has one element and **candidates**$(g_2, \cdot)$ has two elements, or vice-versa. Assume, without loss of generality, that **candidates**$(g_1, \cdot)$ contains one element and **candidates**$(g_2, \cdot)$ contains two elements. Let $u_1$ denote the sole element of **candidates**$(g_1, \cdot)$, and let $v^A \in V(S^O)$ and $u_2 \in V(S^L) \cup V(S^R)$ denote the two elements of **candidates**$(g_2, \cdot)$. Then **candidates**$(g, \cdot) = BLCA_S(u_1, v^A) \cup BLCA_S(u_1, u_2)$. Let $w = lca_S(u_1, u_2)$, $x = llca_S(u_1, u_2)$, and $y = rlca_S(u_1, u_2)$. Now consider the possible membership of $u_1$ and $u_2$ among the nodes of $S$.

If both $u_1$ and $u_2$ are nodes of the same tree $\hat{S} \in \{S^L, S^R\}$, then from fact 1, $w = x = y \in V(\hat{S})$, and so $BLCA_S(u_1, u_2) = \{w\}$. Furthermore, from fact 2, $llca_S(u, v^A) = rlca_S(u, v^A) = v^A$, and so $BLCA_S(u_1, v^A) = \{v^A\}$.

If $u_1$ is a node of $S^O$, then from fact 2, $w = lca_S(u_1, v^A) = x = llca_S(u_1, v^A) = y = rlca_S(u_1, v^A)$. Thus, $BLCA_S(u_1, u_2) = BLCA_S(u_1, v^A) = \{w\}$.

If one of $u_1$ or $u_2$ is a node of $S^L$ and the other is a node of $S^R$, then from fact 3, $w = x = y = v^A$, and so $BLCA_S(u_1, u_2) = \{v^A\}$. Furthermore, from fact 2, $llca_S(u_1, v^A) = rlca(u_1, v^A) = v^A$, and so $BLCA_S(u_1, v^A) = \{v^A\}$.

If $u_1$ is a node of $S^H$ and $u_2$ is a node of $S^L$, then from fact 4a, $w = x \in V(S^L)$ and $y = v^A$, and so $BLCA_S(u_1, u_2) = \{w, v^A\}$. Furthermore, from fact 2, $llca_S(u_1, v^A) = rlca(u_1, v^A) = v^A$, and so $BLCA_S(u_1, v^A) = \{v^A\}$.

If $u_1$ is a node of $S^H$ and $u_2$ is a node of $S^R$, then from fact 4b, $w = y \in V(S^R)$ and $x = v^A$, and so $BLCA_S(u_1, u_2) = \{w, v^A\}$. Furthermore, from fact 2, $llca_S(u_1, v^A) = rlca(u_1, v^A) = v^A$, and so $BLCA_S(u_1, v^A) = \{v^A\}$.

In conclusion, either **candidates**$(g, \cdot) = \{w\}$, or **candidates**$(g, \cdot) = \{v^A\}$, or **candidates**$(g, \cdot) = \{v^A, w\}$, where $w \in V(S^L) \cup V(S^R)$.

**Case 3**: **candidates**$(g_1, \cdot)$ and **candidates**$(g_2, \cdot)$ both have two elements. Let $v^A \in V(S^O)$ and $u_1 \in V(S^L) \cup V(S^R)$ denote the elements of **candidates**$(g_1, \cdot)$, and let $v^A \in V(S^O)$ and $u_2 \in V(S^L) \cup V(S^R)$ denote the elements of **candidates**$(g_2, \cdot)$. Then **candidates**$(g, \cdot) = BLCA_S(v^A, v^A) \cup BLCA_S(u_1, v^A) \cup BLCA_S(u_2, v^A) \cup BLCA_S(u_1, u_2)$.

$BLCA_S(v^A, v^A) = \{v^A\}$.

From fact 2, $llca_S(u_1, v^A) = rlca_s(u_1, v^A) = v^A$ and $llca_S(u_2, v^A) = rlca_S(u_2, v^A) = v^A$, and so, $BLCA_S(u_1, v^A) = BLCA_S(u_2, v^A) = \{v^A\}$.

Let $w = lca_S(u_1, u_2)$, $x = llca_S(u_1, u_2)$, and $y = rlca_S(u_1, u_2)$. If $u_1$ and $u_2$ are nodes of the same tree $\hat{S} \in \{S^L, S^R\}$, then from fact 1, $w = x = y \in V(\hat{S})$, and so $BLCA_S(u_1, u_2) = \{w\}$. If one of $u_1$ or $u_2$ is a node of $S^L$ and the other is a node of $S^R$, then from fact 3, $w = x = y = v^A$, and so $BLCA_S(u_1, u_2) = \{v^A\}$.

In conclusion, either **candidates**$(g, \cdot) = \{v^A\}$, or **candidates**$(g, \cdot) = \{v^A, w\}$, where $w \in V(S^L) \cup V(S^R)$. □

##### S2.2.5 Proof of Theorem 3.7

*Proof.* After a linear-time preprocessing step, LCA operations on a tree can be performed in constant time (Harel and Tarjan 1984, Gabow and Tarjan 1985). By definition, for two nodes $u$ and $v$ of $S$, $BLCA_S(u, v)$ contains $llca_S(u, v) = lca_{S_l}(u, v)$ and $rlca_S(u, v) = lca_{S_r}(u, v)$. Thus, the algorithm can first preprocess $S$ in time $O(|S|)$, after which $BLCA_S(u, v)$ can be computed in constant time.

Recall that Algorithm 1 has complexity is $O(|G| \cdot |S|^5)$ by Theorem 3.4. Compared to Algorithm 1, Algorithm 2 has the following modifications that affect its time complexity:

1. The *for* loop at lines 4-5 requires time $O(|G|)$. [This time complexity is subsumed by the *for* loop at lines 2-3, which remains unchanged and requires time $O(|G| \cdot |S|)$.]

2. The *for* loop at lines 6-8 requires time $O(|G|)$. [This time complexity is unchanged.]

3. At each iteration of the *for* loop at lines 9-22, each of $s_1$ and $s_2$ take on at most two values (by Lemma S2.2), and given $s_1$ and $s_2$, each $s$ takes on at most two values (since $BLCA_S(\cdot,\cdot)$ has at most two elements). Line 21 requires time $O(1)$. [Previously each of $s_1$, $s_2$, and $s$ took on at most $|S|$ values.]

In particular, the third modification has reduced the time complexity by a factor of $O(|S|^3)$, so Algorithm 2 has a time complexity of $O(|G| \cdot |S|^2)$. □

#### S2.3 Allowing Gene Tree Handles

##### S2.3.1 Proof of Lemma 3.8

*Proof.* By Theorem 3.6, line 2 computes $\mathbf{ECrs}(g,s,x)$ correctly for each $g \in V(G_{g_1})$, each $x \in \mathbb{LR}$, and each $s \in \mathbf{candidates}(g,x)$. Since $R_v(gh)$ is restricted to $r(S)$, it suffices to show that $\mathbf{ECrs}(gh, r(S), x)$ is computed correctly for each $x \in \mathbb{LR}$.

Note that line 2 computes $\mathbf{candidates}(g_1,x_1)$ correctly for $x_1 \in \mathbb{LR}$ and computes $\mathbf{ECrs}(g_1,s_1,x_1)$ correctly for $s_1 \in \mathbf{candidates}(g_1,x_1)$. The rest of Algorithm 3 is a straightforward modification of Algorithm 2. Specifically, Algorithm 2 considered an internal gene node $g$ with children $g_1$ and $g_2$ and required a reconciliation $R$ between $G_g$ and $S$ to (a) extend $R^1$ between $G_{g_1}$ and $S$ and extend $R^2$ between $G_{g_2}$ and $S$, and (b) have path $p_1$ between $R_v(g)$ and $R_v(g_1)$ and path $p_2$ between $R_v(g)$ and $R_v(g_2)$. Algorithm 3 instead considers a handle node $gh$ with a single child $g_1$ and requires a reconciliation $R$ between $G_{gh}$ and $S$ to (a) extend $R^1$ between $G_{g_1}$ and $S$ and (b) have path $p_1$ between $R_v(gh) = r(S)$ and $R_v(g_1)$. So, by similar reasoning as in the proofs of Theorem 3.6 and 3.6, the edgeset, signature, and cost of $R$ are computed correctly, and $\mathbf{ECrs}(gh, r(S), x)$ is updated correctly. □

##### S2.3.2 Proof of Lemma 3.9

*Proof.* The time complexity of Algorithm 3 is dominated by the call to ReconcileBLCASimpleSnetwork (line 2), which, by Theorem 3.7, has time complexity $O(|G| \cdot |S|^2)$. □

#### S2.4 Considering Multiple Gene Trees

##### S2.4.1 Definitions over $\mathcal{R}$

$$\begin{aligned}
\mathbf{edgeset}(\mathcal{R}) &= \bigcup_{g \in V(\mathcal{G})} \mathcal{R}_p(g) \\
&= \bigcup_{k=1}^{K} \bigcup_{g \in V(G_k)} R_p^k(g) \\
&= \bigcup_{k=1}^{K} \mathbf{edgeset}(R^k) \\
\mathbf{signature}(\mathcal{R}) &= \mathbf{signature}(\mathbf{edgeset}(\mathcal{R})) \\
&= \mathbf{signature}\left(\bigcup_{k=1}^{K} \mathbf{edgeset}(R^k)\right) \\
&= \sum_{k=1}^{K} \mathbf{signature}(R^k)
\end{aligned}$$

$$
\begin{aligned}
\text{for } e \in E(S), \mathbf{L}_{\mathcal{R}}(e) &= |\{g \in V(\mathcal{G}) : e \in \mathcal{R}_p(g)\}| \\
&= \left| \bigcup_{k=1}^{K} \{g \in V(G_k) : e \in R_p^k(g)\} \right| \\
&= \sum_{k=1}^{K} \left|\{g \in V(G_k) : e \in R_p^k(g)\}\right| \\
&= \sum_{k=1}^{K} \mathbf{L}_{R^k}(e) \\
\mathbf{DC}_{\mathcal{R}} &= \sum_{e \in \mathbf{edgeset}(\mathcal{R})} (\mathbf{L}_{\mathcal{R}}(e) - 1) \\
&= \sum_{e \in \mathbf{edgeset}(\mathcal{R})} \left( \left( \sum_{k=1}^{K} \mathbf{L}_{R^k}(e) \right) - 1 \right)
\end{aligned}
$$

##### S2.4.2 Proof of Lemma 3.10

*Proof.* Let $s = r(S)$ and $x = \mathbf{signature}(\mathcal{Q}) = \mathbf{signature}(\mathcal{R})$. Let $\hat{L}$ be the image of $\bigcup_{k=1}^{K} L(G_k)$ under $Le$, that is, the set of leaves of $S$ that are mapped from the leaves of each tree in $\mathcal{G}$. The result follows as in the proof of Lemma 3.1. □

##### S2.4.3 Proof of Lemma 3.11

*Proof.* The proof uses the same approach as in the proofs of Lemma 3.2 and Lemma 3.3.

Suppose to the contrary that $\mathcal{R}^*$ is an s-optimal reconciliation such that at least one of $\mathcal{R}^{*,k-1}$ or $R^{*,k}$ is not s-optimal. Then, let $\mathcal{R}^{k-1}$ denote an s-optimal reconciliation between $\mathcal{G}^{k-1}$ and $S$ such that $\mathbf{signature}(\mathcal{R}^{k-1}) = \mathbf{signature}(\mathcal{R}^{*,k-1})$. And let $R^k$ denote an s-optimal reconciliation between $G_k$ and $S$ such that $\mathbf{signature}(R^k) = \mathbf{signature}(R^{*,k})$. Consider a new reconciliation $\mathcal{R} = \mathcal{R}^{k-1} \cup \{R^k\}$ between $\mathcal{G}$ and $S$.

**Edgeset of $\mathcal{R}$**

Let $E_1 = \mathbf{edgeset}(\mathcal{R}^{k-1})$ and $E_2 = \mathbf{edgeset}(R^k)$. Then the edgeset of $\mathcal{R}$ is given by

$$
\begin{aligned}
E = \mathbf{edgeset}(\mathcal{R}) &= \bigcup_{u \in V(\mathcal{G})} \mathcal{R}_p(u) \\
&= \left( \bigcup_{u \in V(\mathcal{G}^{k-1})} \mathcal{R}_p(u) \right) \cup \left( \bigcup_{u \in V(G_k)} \mathcal{R}_p(u) \right) \\
&= \left( \bigcup_{u \in V(\mathcal{G}^{k-1})} \mathcal{R}_p^{k-1}(u) \right) \cup \left( \bigcup_{u \in V(G_k)} R_p^k(u) \right) \\
&= \mathbf{edgeset}(\mathcal{R}^{k-1}) \cup \mathbf{edgeset}(R^k) = E_1 \cup E_2.
\end{aligned}
$$

Similarly, let $E_1^* = \mathbf{edgeset}(\mathcal{R}^{*,k-1})$ and $E_2^* = \mathbf{edgeset}(R^{*,k})$. Then the edgeset of $\mathcal{R}^*$ is given by $E^* = E_1^* \cup E_2^*$.

**Cost of $\mathcal{R}$**

Let $c_1 = \mathbf{DC}_{\mathcal{R}^{k-1}}$ and $c_2 = \mathbf{DC}_{R^k}$. Consider the cost of $\mathcal{R}$:

$$
c = \mathbf{DC}_{\mathcal{R}} = \sum_{e \in \mathbf{edgeset}(\mathcal{R})} (\mathbf{L}_{\mathcal{R}}(e) - 1)
$$

Note that for any edge $e \in E$, the following holds:

(1) If $e \in E_1$, then $\mathcal{R}^{k-1}$ contributes $\mathbf{L}_{\mathcal{R}^{k-1}}(e)$ lineages to $\mathbf{L}_{\mathcal{R}}(e)$.

(2) If $e \in E_2$, then $R^k$ contributes $\mathbf{L}_{R^k}(e)$ lineages to $\mathbf{L}_{\mathcal{R}}(e)$.

Then, expanding the cost of $\mathcal{R}$ over the partition of $E$ consisting of sets $E_1 \setminus E_2$, $E_2 \setminus E_1$, and $E_1 \cap E_2$, we get the following:

(i) $$\sum_{e \in E_1 \setminus E_2} (\mathbf{L}_{\mathcal{R}}(e) - 1) = \sum_{e \in E_1 \setminus E_2} (\mathbf{L}_{\mathcal{R}^{k-1}}(e) - 1)$$

(ii) $$\sum_{e \in E_2 \setminus E_1} (\mathbf{L}_{\mathcal{R}}(e) - 1) = \sum_{e \in E_2 \setminus E_1} (\mathbf{L}_{R^k}(e) - 1)$$

(iii) $$\begin{aligned}\sum_{e \in E_1 \cap E_2} (\mathbf{L}_{\mathcal{R}}(e) - 1) &= \sum_{e \in E_1 \cap E_2} (\mathbf{L}_{\mathcal{R}^{k-1}}(e) + \mathbf{L}_{R^k}(e) - 1) \\ &= \sum_{e \in E_1 \cap E_2} (\mathbf{L}_{\mathcal{R}^{k-1}}(e) - 1) + \sum_{e \in E_1 \cap E_2} (\mathbf{L}_{R^k}(e) - 1) + |E_1 \cap E_2|\end{aligned}$$

Combining (i) and the first term of (iii):

$$\begin{aligned}\sum_{e \in E_1 \setminus E_2} (\mathbf{L}_{\mathcal{R}^{k-1}}(e) - 1) + \sum_{e \in E_1 \cap E_2} (\mathbf{L}_{\mathcal{R}^{k-1}}(e) - 1) &= \sum_{e \in E_1} (\mathbf{L}_{\mathcal{R}^{k-1}}(e) - 1) \\ &= \mathbf{DC}_{\mathcal{R}^{k-1}} = c_1\end{aligned}$$

Similarly, combining (ii) and the second term of (iii):

$$\begin{aligned}\sum_{e \in E_2 \setminus E_1} (\mathbf{L}_{R^k}(e) - 1) + \sum_{e \in E_1 \cap E_2} (\mathbf{L}_{R^k}(e) - 1) &= \sum_{e \in E_2} (\mathbf{L}_{R^k}(e) - 1) \\ &= \mathbf{DC}_{R^k} = c_2\end{aligned}$$

Thus, after some rearrangement, the cost of $\mathcal{R}$ is given by

$$c = c_1 + c_2 + \left|E_1 \cap E_2\right|.$$

Similarly, let $c_1^* = \mathbf{DC}_{\mathcal{R}^{*,k-1}}$ and $c_2^* = \mathbf{DC}_{R^{*,k}}$. Then the cost of $\mathcal{R}^*$ is given by $c^* = c_1^* + c_2^* + \left|E_1^* \cap E_2^*\right|$.

By Lemma 3.10, $E_1 = E_1^*$ and $E_2 = E_2^*$. By assumption, at least one of $\mathcal{R}^{*,k-1}$ or $R^{*,k}$ is not s-optimal, and $\mathcal{R}^{k-1}$ and $R^k$ are s-optimal, so it follows that $c_1^* + c_2^* > c_1 + c_2$. Thus,

$$c^* = c_1^* + c_2^* + \left|E_1^* \cap E_2^*\right| > c_1 + c_2 + \left|E_1 \cap E_2\right| = c,$$

which contradicts the s-optimality of $\mathcal{R}^*$. □

###### S2.4.4 Proof of Lemma 3.12

*Proof.* The proof uses the same approach as in the proof of Lemma 3.3 and Lemma 3.6. It suffices to show that for each $k$ such that $1 \le k \le K$ and each $x \in \mathbb{LR}$, the value of $\mathbf{ECs}(G_k, x)$ is computed correctly. Our proof is by induction on $k$ (i.e. by induction on $G_k$ in the set $\{G_1, \ldots, G_K\}$).

**Base Case**

If $k = 1$, then $\mathcal{G}^1 = \{G_1\}$. Given a signature $x \in \mathbb{LR}$, an s-optimal reconciliation $R^1$ between $\mathcal{G}^1$ and $S$ such that $R_v^1(gh_1) = r(S)$ and $\mathbf{signature}(R^1) = x$ must be rs-optimal. By Lemma 3.8, the edgeset and cost of $R^1$ are found in $\mathbf{ECrs}(gh_1, r(S), x)$. These entries are correctly stored in $\mathbf{ECs}(G_1, x)$ after the execution of the *for* loop in lines 7-8.

**Inductive Case**
Let $k > 1$. Let us assume that entries $\mathbf{ECs}(G_{k-1}, x)$ are computed correctly for each $x \in \mathbb{LR}$. Based on this induction hypothesis, we will show that the entries $\mathbf{ECs}(G_k, x)$ are computed correctly for each $x \in \mathbb{LR}$.

Let $\mathcal{R}^{k-1} = (\mathcal{R}_v^{k-1}, \mathcal{R}_p^{k-1})$ be an s-optimal reconciliation between $\mathcal{G}^{k-1}$ and $S$ such that $\mathbf{signature}(\mathcal{R}^{k-1}) = x_1$. Then $\mathcal{R}^{k-1}$ has edgeset $E_1$ and cost $c_1$ given by $\mathbf{ECs}(G_{k-1}, x)$. Similarly, let $R^k$ be an s-optimal reconciliation between $G_k$ and $S$ with an analogous edgeset $E_2$ and cost $c_2$ given by $\mathbf{ECrs}(gh_k, r(S), x_2)$. Now consider a reconciliation $\mathcal{R} = \mathcal{R}^{k-1} \cup \{R^k\}$ between $\mathcal{G}^k$ and $S$.

Let $E$ and $c$ denote the edgeset and cost of $\mathcal{R}$. We first show that the edgeset $E$, signature $x$, and cost $c$ of $\mathcal{R}$ are computed correctly, then show that $\mathbf{ECs}(G_k, x)$ stores the edgeset and cost for some reconciliation that is s-optimal.

**Edgeset of $\mathcal{R}$**
As in the proof of Lemma 3.11, the edgeset of $\mathcal{R}$ is given by $E = E_1 \cup E_2$, as computed in line 13.

**Signature of $\mathcal{R}$**
The signature of $\mathcal{R}$ is given by

$$\begin{aligned} x &= \mathbf{signature}(\mathcal{R}) = \mathbf{signature}(E) \\ &= \mathbf{signature}(E_1 \cup E_2) \\ &= \mathbf{signature}(E_1) + \mathbf{signature}(E_2) \\ &= \mathbf{signature}(R_1) + \mathbf{signature}(R_2) \\ &= x_1 + x_2, \end{aligned}$$

as computed in line 15.

**Cost of $\mathcal{R}$**
Finally, consider the cost of $\mathcal{R}$. As in the proof of Lemma 3.11,

$$c = c_1 + c_2 + |E_1 \cap E_2|,$$

as computed in line 14.

**Update of $\mathbf{ECs}(G_k, x)$**
Lastly, note that the *for* loop at line 10 searches over all possible signatures $x_1 \in \mathbb{LR}$ for $\mathcal{R}^{k-1}$ and $x_2 \in \mathbb{LR}$ for $R^k$. Thus, at the end of the *for* loop, the algorithm will have evaluated the edgeset $E$, signature $x$, and cost $c$ for every reconciliation $\mathcal{R}^k$ subject to the previously stated constraints. To update $\mathbf{ECs}(G_k, x)$, what remains is to retain only the edgeset and cost for some reconciliation that is s-optimal with respect to signature $x$. This filter is executed in lines 16-17.

**Minimum Reconciliation Cost**
Finally, the minimum reconciliation cost between $\mathcal{G}^K$ and $S$ is simply $\min_{x \in \mathbb{LR}} \mathbf{cost}(\mathbf{ECs}(G_K, x))$, as computed in line 18. □

##### S2.4.5 Proof of Lemma 3.13

*Proof.* Consider the time complexity of Algorithm 4 step-by-step. The *for* loop at lines 2-3 requires time $O(K)$. The *for* loop at lines 4-17 is performed $K$ times, at each iteration processing tree $G_k$. By Lemma 3.9, each call to ReconcileWithHandleSimpleSnetwork (line 5) takes time $O(|G_k| \cdot |S|^2)$. If $k = 1$, the *for* loop at lines 7-8 takes time $O(1)$. Otherwise, each of $x_1$ and $x_2$ take on four values (line 10). As in Algorithm 1 and Algorithm 2, lines 13 and 14 each require at most time $O(|S|^2)$, and each other statement in lines 11-17 requires time $O(1)$. Therefore, lines 4-17 require time $O(\sum_{G_k \in \mathcal{G}} |G_k| \cdot |S|^2)$. Finally, computing the minimum in line 18 over a set of size four takes time $O(1)$. □

#### S2.5 Putting the Pieces Together

##### S2.5.1 Additional Algorithms

**Algorithm S1**

**function** RECONCILELCASTREE($G$, $S$, $Le$)
**input** gene tree $G$, species tree $S$, leaf map $Le$
**output** mapping **candidates**$(g)$, mapping **ECrs**$(g, s)$
  **for** each $g \in L(G)$ **do**
    Set **ECrs**$(g, Le(g)) = (\emptyset, 0)$.
    Set **candidates**$(g) = \{Le(g)\}$.
  **for** each $g \in I(G)$ in post-order **do**
    Set $(g_1, g_2) = c(g)$.
    Set $s_1$ and $s_2$ to be the single elements in **candidates**$(g_1)$ and **candidates**$(g_2)$, resp.
    Set $s = lca_S(s_1, s_2)$.
    Set $p_1$ and $p_2$ to be the single elements in $paths_S(s, s_1)$ and $paths_S(s, s_2)$, resp.
    Set $(E_1, c_1) =$ **ECrs**$(g_1, s_1)$.
    Set $(E_2, c_2) =$ **ECrs**$(g_2, s_2)$.
    Set $E = E_1 \cup E_2 \cup p_1 \cup p_2$.
    Set $c = c_1 + c_2 + |E_1 \cap E_2| + |E_1 \cap p_2| + |E_2 \cap p_1| + |p_1 \cap p_2|$.
    Set **candidates**$(g) = \{s\}$.
    Set **ECrs**$(g, s) = (E, c)$.
  **return candidates**, **ECrs**.

**Algorithm S2**

**function** RECONCILEWITHHANDLESTREE($G$, $S$, $Le$)
**input** gene tree $G$ with handle $(gh, g_1)$, species tree $S$, leaf map $Le$
**output** mapping **ECrs**$(g, s, x)$
  Set **candidates**, **ECrs** = RECONCILELCASTREE$(G_{g_1}, S, Le)$.
  Set $s_1$ to be the single element in **candidates**$(g_1)$.
  Set $p_1$ to be the single element in $paths_S(r(S), s_1)$.
  Set $(E_1, c_1) =$ **ECrs**$(g_1, s_1)$.
  Set $E = E_1 \cup p_1$.
  Set $c = c_1$.
  Set **ECrs**$(gh, r(S)) = (E, c)$.
  **return ECrs**.

**Algorithm S3**

**function** ReconcileMultipleStree($\mathcal{G}$, $S$, $Le$)
**input** forest $\mathcal{G}$ of gene trees $\{G_1, \ldots, G_K\}$ with handle nodes $\{gh_1, \ldots, gh_K\}$, species tree $S$, leaf map $Le$
**output** minimum reconciliation cost between $\{G_1, \ldots, G_K\}$ and $S$ such that each node of $\{gh_1, \ldots, gh_K\}$ is mapped to $r(S)$
  **for** each $k$ from 1 to $K$ **do**
    Set $\mathbf{ECrs} = $ ReconcileWithHandleStree($G_k, S, Le$).
    **if** $k = 1$ **then**
      Set $\mathbf{ECs}(G_1) = \mathbf{ECrs}(gh_1, r(S))$.
    **else**
      Set $(E_1, c_1) = \mathbf{ECs}(G_{k-1})$.
      Set $(E_2, c_2) = \mathbf{ECrs}(gh_k, r(S))$.
      Set $E = E_1 \cup E_2$.
      Set $c = c_1 + c_2 + |E_1 \cap E_2|$.
      Set $\mathbf{ECs}(G_k) = (E, c)$.
  **return cost**($\mathbf{ECs}(G_K)$).

##### S2.5.2 Proof of Lemma 3.16

*Proof.* The proof is analogous to that of Lemma 3.5.

Let $g_1$ and $g_2$ denote the children of $g$. Let $s = R_v(g)$, $s_1 = R_v(g_1)$, and $s_2 = R_v(g_2)$, and let $p = R_p(g)$, $p_1 = R_p(g_1)$, and $p_2 = R_p(g_2)$. Let $M = \mathcal{M}(s)$, $M_1 = \mathcal{M}(s_1)$, and $M_2 = \mathcal{M}(s_2)$, and let $B = \mathcal{B}(g)$, $B_1 = \mathcal{B}(g_1)$, and $B_2 = \mathcal{B}(g_2)$.

By Definition 2.1, for each $v \in L(G_g)$, $Le(v) \leq_S R_v(g)$. By Definition 3.4, $B$ is the lowest biconnected component of $S$ such that $L(S_{r(B)})$ contains $\{Le(v) \mid v \in L(G_g)\}$, so it follows that $B \leq_S M$. By assumption, $M \neq B$, so it follows that $B <_S M$. By assumption, $M_1 = B_1$ and $M_2 = B_2$. By Definition 3.4, it is also easily shown that $\mathring{B} = lca_{bc(S)}(\mathring{B}_1, \mathring{B}_2) = lca_{bc(S)}(\mathring{M}_1, \mathring{M}_2)$. Since $\mathring{B} = lca_{bc(S)}(\mathring{M}_1, \mathring{M}_2)$ and $\mathring{B} <_{bc(S)} \mathring{M}$, any path from $\mathring{M}$ to $\mathring{M}_1$ must contain $\mathring{B}$, and any path from $\mathring{M}$ to $\mathring{M}_2$, must contain $\mathring{B}$.

Let $s^* = r(B)$ so that $\mathcal{M}(s^*) = B$. Since $s \in V(M)$ and $B <_S M$, $s^* <_S s$. By Definition 2.1, $p_1$ is a path from $s$ to $s_1$. Obtain $\mathring{p}_1$ from $p_1$ by replacing each node $u$ in $p_1$ with $\mathring{\mathcal{M}}(u)$ and removing duplicate nodes. Then $\mathring{p}_1$ is a path from $\mathring{M}$ to $\mathring{M}_1$ and so contains $\mathring{B}$. Thus, $p_1$ contains some node $u_1 \in V(B)$. Since $p_1$ starts from $s \notin V(B)$ and contains some node $u_1 \in V(B)$, it is easily shown that $p_1$ must contain $r(B)$. Thus, $p_1$ contains $s^*$, and similarly, $p_2$ contains $s^*$.

Consider the following reconciliation $R^* = (R^*_v, R^*_p)$:

$$R^*_v(u) = \begin{cases} s^*, & \text{if } u = g \\ R_v(u), & \text{otherwise} \end{cases}$$

$$R^*_p(u) = \begin{cases} \emptyset, & \text{if } u = g \text{ and } u = r(G) \\ p \text{ extended with the edges of } p_1 \text{ from } s \text{ to } s^*, & \text{if } u = g \text{ and } g \neq r(G) \\ p_1 \text{ truncated to start at } s^*, & \text{if } u = g_1 \\ p_2 \text{ truncated to start at } s^*, & \text{if } u = g_2 \\ R_p(u), & \text{otherwise} \end{cases}$$

Note that $R^*$ is a valid reconciliation. Then, as in the proof of Lemma 3.5, $R^*$ subsumes $R$. □

##### S2.5.3 Proof of Theorem 3.17

*Proof.* Let $R$ denote an optimal reconciliation between $G$ and $S$. By Corollary 3.16.1, the algorithm need only consider reconciliations $R$ that are consistent with $\mathcal{B}$ and $\mathcal{M}$; that is, for each $g$ of $G$, $R$ must satisfy $\mathcal{M}(R_v(g)) = \mathcal{B}(g)$.

Let $\{B_1, \ldots, B_p\}$ denote the biconnected components of $S$ that are not leaf nodes. For each $i$ such that $1 \le i \le p$, let $R^i$ denote the reconciliation $R$ restricted to $\mathcal{G}_{B_i}$. We will show that $R^i$ corresponds to a reconciliation between $\mathcal{G}_{B_i}$ and $S(B_i)$ with some leaf map $Le_{B_i}$; that is, when computing $R^i$, it suffices to consider $\mathcal{G}_{B_i}$ in place of $G$, $S(B_i)$ in place of $S$, and $Le_{B_i}$ in place of $Le$.

Consider a biconnected component $B_i$. By Corollary 3.16.1 and the fact that $R$ extends $R^i$, for each tree $H \in \mathcal{G}_{B_i}$ and each node $u \in I(H)$, it follows that $\mathcal{M}(R_v(u)) = \mathcal{B}(u) = B_i$. By Lemma 3.15, for each tree $H \in \mathcal{G}_{B_i}$ and each leaf node $v \in L(H)$, $\mathcal{B}(v)$ is a child of $B_i$. Then the set of handle nodes of $\mathcal{B}(v)$ is equal to the set of leaf nodes of $B_i$, so it follows that $R_v(v) = r(\mathcal{B}(v))$ . Furthermore, $r(\mathcal{B}(v))$ is a leaf of $S(B_i)$, so it follows that for each node $u$ of $\mathcal{G}_{B_i}$, $R_v(u) \in V(S(B_i))$ . Thus, for $g \in L(\mathcal{G}_{B_i})$, $Le_{B_i}(g) = r(\mathcal{B}(g))$, as computed in line 8, and $R^i$ is a forest reconciliation between $\mathcal{G}_{B_i}$ and $S(B_i)$, whose minimum cost can be computed as in lines 10 and 12.

Next, it suffices to show that $\mathbf{DC}_R = \sum_{i=1}^{p} \mathbf{DC}_{R^i}$. If so, then to minimize $\mathbf{DC}_R$, it suffices to minimize each $\mathbf{DC}_{R^i}$, and furthermore, the costs of each reconciliation $R^i$ can be added as in line 13. Note that the networks $\{S(B_1), \ldots, S(B_p)\}$ are disjoint, and each edge of $E(S)$ is contained in exactly one $E(S(B_i))$. Thus,

$$\mathbf{DC}_R = \sum_{e \in E(S)} \mathbf{XL}_R(e) = \sum_{i=1}^{p} \left( \sum_{e \in E(S(B_i))} \mathbf{XL}_{R^i}(e) \right) = \sum_{i=1}^{p} \mathbf{DC}_{R^i} \,.$$

□

##### S2.5.4 Proof of Theorem 3.18

*Proof.* Consider the time complexity of Algorithm 5 step-by-step. Let $p$ denote the number of biconnected components of $S$. To and Scornavacca (2015) showed that all biconnected components of $S$ and $M(\cdot)$ can be computed in time $O(|S|)$, $B(\cdot)$ can be computed in time $O(|G_S|)$, $G_S$ can be constructed in total time $O(|G_S| + |S|)$, $\mathcal{G}_{B_i}$ for all $B_i$ can be constructed in time $O(|G_S|)$, and in the worst case, $O(|G_S|) = O(|G| \cdot p)$ (Proof of Theorem 2). Note that our revised definitions of $G_S$ and $\mathcal{G}_{B_i}$ do not change these complexities. Thus, the computation in lines 2, 3, and 4 require time $O(|G| \cdot p + |S|)$.

The *for* loop at lines 6-13 considers each biconnected component $B_i$ of $S$ that is not a leaf. For each $B_i$, the *for* loop at lines 7-8 requires time $O(|\mathcal{G}_{B_i}|)$. Next, the algorithm calls either ReconcileMultipleStree (line 10) or ReconcileMultipleSnetwork (line 12), which, by Theorem 3.13, have time complexity $O(|\mathcal{G}_{B_i}| \cdot |S(B_i)|^2)$. Thus, lines 6-8 require time $O(\sum_{i=1}^{p} |\mathcal{G}_{B_i}| \cdot |S(B_i)|^2)$. Since $|\mathcal{G}_{B_i}| \le |G|$ and $\sum_{i=1}^{p} |S(B_i)|^2 \le |S|^2$, lines 6-8 require time $O(|G| \cdot |S|^2)$, which subsumes $O(|G| \cdot p + |S|)$. □
